# Hybridization kinetics of out-of-equilibrium mixtures of short RNA oligonucleotides

**DOI:** 10.1101/2022.06.09.495531

**Authors:** Marco Todisco, Jack W. Szostak

## Abstract

Hybridization and strand displacement kinetics determine the evolution of the base-paired configurations of mixtures of oligonucleotides over time. Although much attention has been focused on the thermodynamics of DNA and RNA base pairing in the scientific literature, much less work has been done on the time dependence of interactions involving multiple strands, especially in RNA. Here we provide a study of oligoribonucleotide interaction kinetics and show that it is possible to calculate the association, dissociation and strand displacement rates displayed by short oligonucleotides (5nt – 12nt) as simple functions of oligonucleotide length, CG content, ΔG of hybridization and ΔG of toehold binding. We then show that the resultant calculated kinetic parameters are consistent with the experimentally observed time dependent changes in concentrations of the different species present in mixtures of multiple competing RNA strands. We show that by changing the mixture composition, it is possible to create and tune kinetic traps that extend by orders of magnitude the typical sub-second hybridization timescale of two complementary oligonucleotides. We suggest that the slow equilibration of complex oligonucleotide mixtures may have facilitated the nonenzymatic replication of RNA during the origin of life.

## INTRODUCTION

Characterizing the behavior of mixtures of nucleic acids is important in a wide variety of fields, including gene silencing (1-3), primer design for routine PCR experiments, DNA nanotechnology (4,5), the development of novel materials (6-8), and the study of RNA World scenarios relevant to understanding the origins of life (9-12). The pillar on which all these studies have been built is the nearest-neighbor thermodynamic model for the stability of base-paired nucleic acid duplexes (13,14). This model accurately predicts the change in free energy (ΔG°) when two oligonucleotides anneal, at least in cases where the process can be approximated as a two-state transition and assuming that the oligonucleotides have negligible secondary structure and self-interactions.

Although the nearest-neighbor model provides a good description of the equilibrium state of the system, a solution of many different oligonucleotides may require a long time to equilibrate upon mixing, because the system can become trapped in local energy minima. In such cases escaping from metastable states may require thermal annealing to reach the global minimum. Depending on the system, such out-of-equilibrium states can either constitute a strong disadvantage, causing the formation of unwanted duplexes whose concentration varies with time, or it can be a valuable source of complexes with single-stranded overhangs or gaps that can behave as substrates for desired physical processes (15,16), or for chemical reactions such as template copying or loop closing under the proper activating conditions (11,17-19). While the change in free energy can ultimately be calculated with a reasonable degree of accuracy from oligonucleotide sequences and ionic conditions, a more rigorous and extensive study of hybridization kinetics together with modelling is required to better understand the out-of-equilibrium behavior of mixtures of RNA oligonucleotides.

When two complementary oligonucleotides are mixed in solution, they rapidly hybridize to produce a double helix, annealing *via* a two-step process involving nucleation by the formation of several consecutive paired bases, followed by a rapid zippering step. Because the nucleation step is generally assumed to be rate-limiting, duplex formation can be modeled as a bimolecular reaction with an effective rate *k*_*on*_, dominated by the time necessary for the two strands to collide and then form a productive nucleation region that will zipper (20-22). The kinetics of oligonucleotide binding has traditionally been measured using techniques such as following the FRET signal between fluorescently labeled oligonucleotides (1,23,24), following changes in UV or IR absorbance (25-29) or using specific sensors (30-33) that rely on DNA binding to a functionalized surface.

With the aim of better understanding nucleic acid hybridization, a series of modeling approaches have been explored in the past, from approximate analytical treatments (27,34) to Markov-chain stochastic simulations (35) and molecular dynamics coarse-grained simulations (20,36). Recently two noteworthy approaches have been established for predicting the hybridization kinetics of nucleic acids: (i) Zhang and colleagues published a tool based on a complex Weighted Neighbor Voting algorithm built on a database of 36nt long DNA oligonucleotides (37), while (ii) Rejali and colleagues attempted to deconstruct the hybridization rate of a set of short DNA strands with a NN parameterization. Unfortunately, this latter approach showed a limited reliability and, to overcome this issue, the authors proposed a simplified predictive function based on the CG content alone of the analyzed strands. One caveat of these predictive approaches is that they neglect the potential role of oligonucleotide length in association kinetics. The role of length remains controversial in the literature: early studies on long DNA molecules have clearly shown a positive dependence of *k*_*on*_ on the length of the oligonucleotides (34,38). However, the behavior of short oligonucleotides is still a matter of debate, with recent studies claiming to show hybridization rates being either positively dependent, independent or negatively dependent on length (20,22,23,25,27). To further complicate the picture of binding kinetics, studies on DNA and RNA by Cisse at al. have shown that a minimum of 7 consecutive complementary nucleotides are necessary for fast annealing, with a significant drop in rate for shorter stretches (1). This observation suggests that the minimum nucleation site could be larger than previously expected and could thus dramatically influence *k*_*on*_ for the short DNA and RNA oligonucleotides that are the primary focus of our work.

Following their hybridization, two oligonucleotides are not irreversibly bound; instead, they will dissociate with a characteristic dissociation rate *k*_*off*_. While literature *k*_*on*_ values for short DNA and RNA molecules do not typically vary by more than an order of magnitude across different oligonucleotide compositions and lengths, reported duplex lifetimes show a strong dependence on the length and thermodynamic stability of the oligonucleotide duplexes, resulting in *k*_*off*_ values spanning many orders of magnitude (23,27,39).

When more than two complementary sequences are present in solution, individual association and dissociation rates are not sufficient to provide a complete description of the time-dependent evolution of the system. An important phenomenon that must be accounted for is toehold-mediated strand displacement. This occurs when one strand of a duplex is displaced by an incoming invading strand, a phenomenon that is widely exploited in DNA nanotechnology to develop complex circuits (40-42). More recently toehold-mediated strand displacement has been applied in the field of the origin of life to enable enzyme-free copying of duplex RNA (43). A seminal study by Zhang and Winfree (44) has clarified the details of this process in DNA and shown how the magnitude of the strand displacement rate depends on the binding of the incoming oligonucleotide to an unpaired region of the template, the so-called toehold. Only recently have the kinetics of toehold-mediated strand displacement for RNA been computationally (45) and experimentally investigated, using an approach analogous to that of the Winfree group by relying on a fluorescent reporter to monitor strand displacement over a region longer than 20bp (46). However, further studies are needed to provide a deeper understanding of this process in shorter oligonucleotides and establish a reliable and comprehensive model for the prediction of rates of toehold-mediated strand displacement in RNA oligonucleotides.

Together, *k*_*on*_, *k*_*off*_ and toehold-mediated strand displacement rates should enable the description of the evolution of a mixture of oligonucleotides from their mixing to their equilibrium state, giving valuable information on the state of the system at intermediate times. In this work we present a study of RNA annealing kinetics and provide a framework for the prediction of the time-dependent evolution of mixtures of oligoribonucleotides.

## MATERIALS AND METHODS

### General

Controlled pore glass (CPG) columns, phosphoramidites and reagents for oligonucleotide synthesis and purification were from Glen Research. Reagents for cleavage and deprotection were from Sigma-Aldrich. Buffers were prepared from 1 M Tris stock solutions (Invitrogen) and pH-adjusted using HCl. Sodium chloride and magnesium chloride hexahydrate were from Sigma-Aldrich. Unless otherwise stated all measurements were performed in 200 mM Tris-HCl pH 8 and 100 mM MgCl_2_. Because the literature standard for nucleic acids is 1 M NaCl, we performed a subset of experiments in 5 mM Tris-HCl 1 M NaCl pH 7 to compare the two conditions and no significant difference was noted, and so we consider these two conditions as equivalent (see Supplementary Data 1 and 5).

### Solid phase synthesis of oligoribonucleotides

RNA oligonucleotides were synthesized in-house on an Expedite 8909 by solid-phase synthesis using the DMT-on protocol. Synthesized oligonucleotides were cleaved from the RNA-CPG columns with AMA (1.2 ml, 1:1 mixture of 40% methylamine solution and 28% NH_4_OH) for 30 minutes. The solution containing the oligonucleotides was transferred from the column to a screw-cap vial and incubated at 65°C for 20 minutes for deprotection. The resulting solution was dried for 30 minutes at 40°C in a speed-vac and then lyophilized overnight. To remove the TBDMS protecting groups the resultant powder was resuspended in DMSO (115 ul), TEA (60 ul) and TEA^.^3HF (75 ul) and incubated at 65°C for 2.5 hours. Oligonucleotides were purified from the mixture and the DMT group removed using GlenPak columns.

Oligonucleotide concentrations were determined spectrophotometrically using a NanoDrop 2000 (Thermo Scientific) using the extinction coefficients computed according to Tataurov et al. (47).

We monitored the extent of hybridization of one oligonucleotide by measuring the fluorescent emission of 2-aminopurine (2Ap), a fluorescent adenine base analogue whose quantum yield diminishes when in a double helix. In the absence of published values for the extinction coefficients of 2Ap-containing RNA strands, we found that the extinction coefficients provided for 2Ap-DNA by Xu and Nordlund (48) and by the IDT oligo analyzer tool closely approximate the values of 2Ap-RNA, validated by us through quantitative quenching experiments against complementary strands of known concentration (see Supplementary Data 2).

### Measurement of binding affinity through melting experiments

Although there is a wide body of literature on computing the free energy of binding and thus the affinity of complementary oligonucleotides, we began by verifying the predictions of the nearest neighbor-model in our specific experimental system, in order to exploit the model and the crucial relationship between *K*_*D*_ and binding and dissociation kinetics (*K*_*D*_ = *k*_*off*_*/k*_*on*_).

We measured the binding energies of highly stable oligonucleotide duplexes (ΔG < -10kcal/mol) through melting experiments monitored using either a Jasco FP-8500 Spectrofluorometer equipped with ETC-815 Peltier temperature-controlled cell holder (for fluorescently labeled 2-aminopurine containing oligonucleotides) or an Agilent 3500 UV-Vis Spectrophotometer (for non-fluorescent oligonucleotides). For each pair of oligonucleotides, we measured melting curves using a temperature ramp of 3°C/min. All curves acquired this way have been corrected for baselines at low and high temperature and normalized from zero to one to determine the fraction of unbound oligonucleotides at every temperature. One of the advantages of measuring the thermal stability of a duplex by fluorescence over conventional UV absorbance, is that we can use a complementary non-fluorescent oligonucleotide at a very high concentration while maintaining a high signal to background ratio. Studying the mixture of a 2Ap-containing oligonucleotide (A) and a binding non-fluorescent oligonucleotide (B), having concentrations respectively [A] and [B], with [B] = [A] + Δ, we can track the fluorescent signal resulting from the melting of the duplex and the release of A during a temperature ramp. The normalized amplitude of the fluorescent trace reflects the fraction (f) of unbound fluorescent oligonucleotide [A] over its total concentration [A_tot_].

We can write the dissociation constant K_D_ for the binding of A and B as follows:

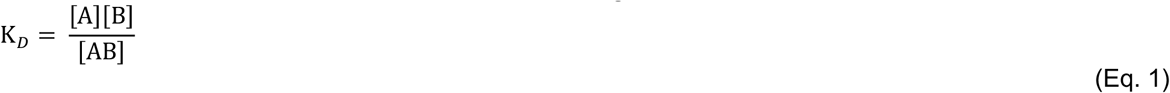

We can substitute concentrations, [B] = [A] + Δ and [AB] = [A_tot_] - [A]:

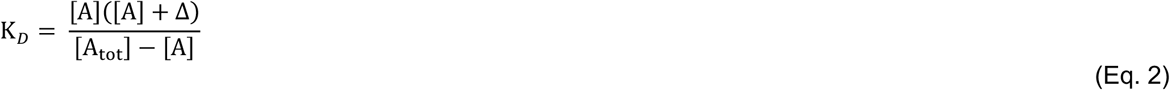

And then substitute [A] as f · [A_tot_] and rearrange:

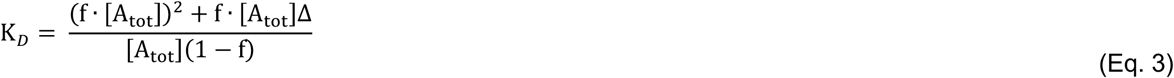

Using this expression, we can readily calculate K_D_ at every temperature of our melting curves since [A_tot_] and Δ are known by preparation. A Van’t Hoff plot for K_D_ vs 1/T can then be used to obtain ΔH and ΔS according to the following function:

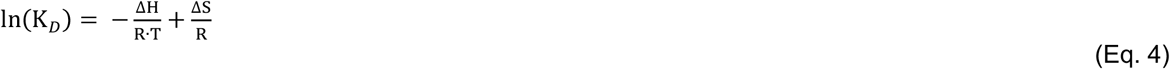

For UV melting experiments, the two oligonucleotides were introduced at equal concentration to maximize the signal to noise ratio. In this condition the mathematical description simplifies and the thermodynamic parameters are extracted from a Van’t Hoff plot of ln(c_tot_/4) vs 1/Tm, where c_tot_/4 is the dissociation constant at the Tm and Tm is the melting temperature of the duplex (the temperature at which the fraction of duplexes in solution is equal to 0.5) in Kelvin.

### Stopped-Flow measurements of association kinetics

For fluorescence measurements we find a lower bound for useful signal detection of 2-aminopurine at ≈ 10 nM using a 1 cm path-length quartz cuvette. Because *k*_*on*_ for nucleic acids at high ionic strength is typically larger than 10^6^ M^-1^s^-1^, reactions at accessible concentrations reach equilibrium in a matter of seconds and are thus too fast to be properly studied by hand mixing. The time dependence of binding was therefore measured using a Jasco FP-8500 Spectrofluorometer equipped with the SFS-852T Stopped-Flow accessory, using samples with concentrations in the micromolar range. To convert fluorescent signals to concentrations, all collected data were normalized using a control experiment where the fluorescent oligonucleotide was mixed only with buffer. Data processed in this way were fit in MATLAB as a simple bimolecular reaction modeled through a set of numerically integrated differential equations. For weakly binding oligonucleotides, *k*_*off*_ and *k*_*on*_ were globally fit over a series of measurements at different concentrations (see Supplementary Data 3 for detailed description of fitting procedure).

### Measurement of strand displacement

If the binding of an oligonucleotide to a toehold region equilibrates on a time scale much faster than the subsequent strand displacement reaction, which is likely for the short toehold lengths explored in this work, we can approximate the strand displacement process as a simple second order reaction with an effective second order rate constant *k*_*displ*_ *= k*_*s*_*⋅K*_*A*_, where *k*_*s*_ is the rate for the unimolecular process of branch migration and *K*_*A*_ is the association constant of the toehold with the incoming invading oligonucleotide (*K*_*A*_ *= K*_*D*_^*-1*^).

To measure strand displacement, we exploited the properties of 2-aminopurine to track either (i) the release of a bound oligonucleotide bearing the fluorescent moiety as an increase in fluorescence emission over time, (ii) the quenching of a 2-aminopurine nucleotide located in the toehold region of the templating oligonucleotide, or (iii) the quenching of 2-aminopurine located in the incoming strand-displacing oligonucleotide. Data collected in this way for mixtures at various concentrations of displacing oligonucleotide were fit as second order irreversible reactions with rate *k*_*displ*_.

### Measurement of the time-dependent evolution of oligonucleotide mixtures

To measure the equilibration process in a mixture of oligonucleotides, we prepared solutions of oligonucleotides competing for shared binding sites. For a typical experiment four sequences A_S_, A_L_, B_S_, B_L_ of short (S) or long (L) length are mixed. The A and B sequences are complementary, so that every A sequence can interact with every B sequence while self-interactions are negligible at our experimental concentrations. To track the evolution of the system, we strategically placed 2-aminopurine in B_L_ so that it could be quenched only by A_L_. For every experiment two solutions, one containing A_S_ and A_L_ and one containing B_S_ and B_L_ were mixed and 2-aminopurine fluorescence was monitored over time. When the four oligonucleotides are initially mixed (i.e. at t_0_) we observe the fluorescent signal coming from B_L_ (see Figure 1), but within a few seconds only double helices A_L_B_L_, A_S_B_S_, A_L_B_S_ and A_S_B_L_ are present and the residual signal from 2Ap comes only from A_S_B_L_ duplexes leading to partial quenching of the 2Ap signal. Over time the system converges to its energy minimum where only A_L_B_L_ and A_S_B_S_ duplexes are present and all fluorescence is quenched. The rate of fluorescence decay can therefore be treated as a result of the complex interplay of *k*_*on*_, *k*_*off*_ and *k*_*displ*_, as the system converges to its energy minimum.

**Figure 1.**
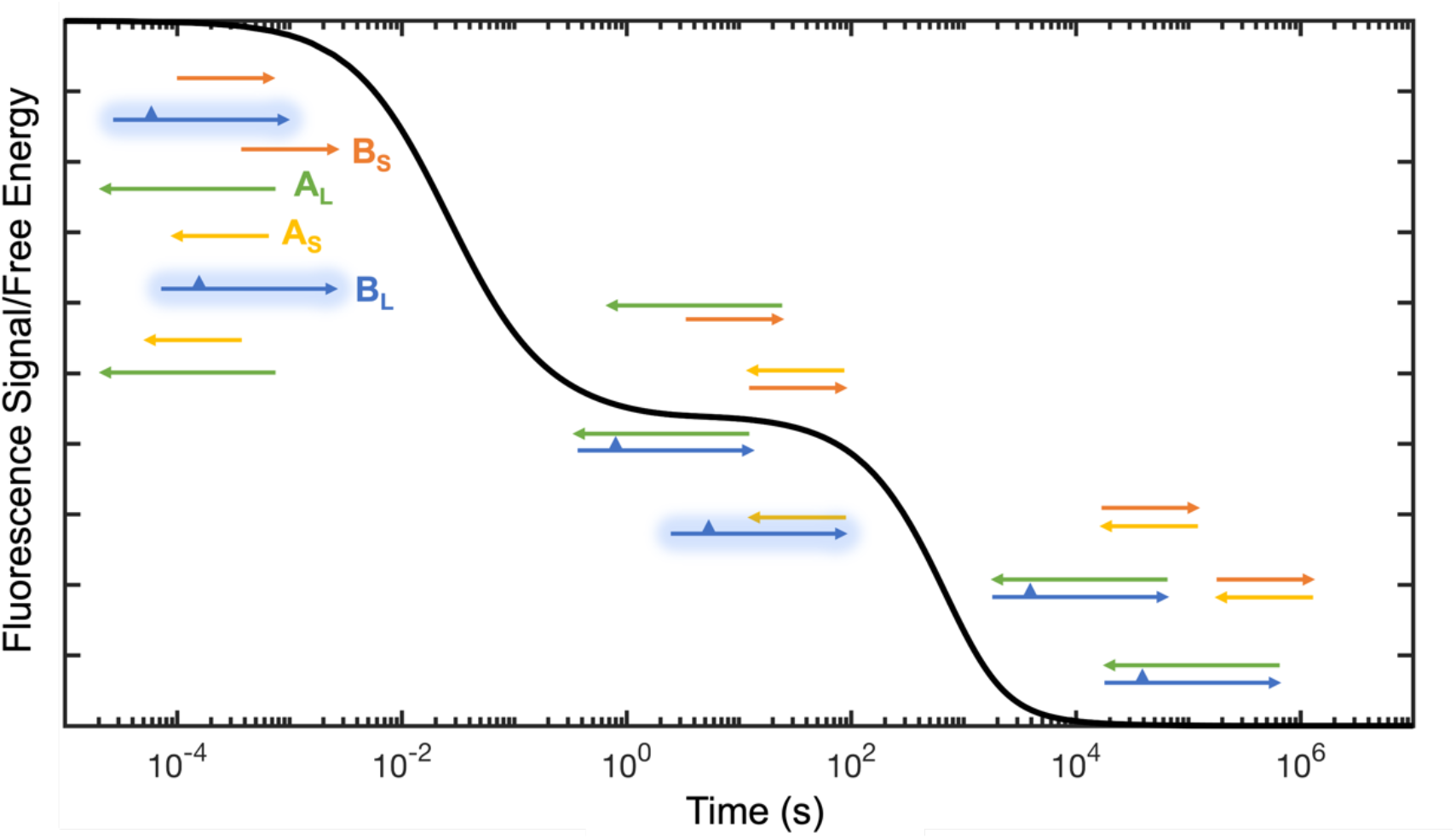
Experimental design for measurement of equilibration time in mixtures of oligonucleotides. At the beginning of the experiment the fluorescent signal of 2Ap (depicted as a triangle) coming from oligonucleotide B_L_ is maximal, and is progressively quenched as more A_L_B_L_ duplex is formed. A curve depicting a hypothetical experiment is shown here in black.

## RESULTS

### Experimental Design

Although fluorescence-based detection of annealing is advantageous because of high sensitivity and easy accessibility, the conjugation of dyes to oligonucleotides can change their properties (49-51). To avoid altering the properties of the oligonucleotides while also retaining the advantages of fluorescence, we use 2-aminopurine (2Ap) as an internal reporter of the state of the system. 2Ap is a fluorescent adenine analogue with an excitation maximum at ≈ 305 nm and an emission maximum at ≈ 370 nm. The quantum yield of 2Ap is diminished in a stacked conformation and therefore its fluorescence emission varies according to the extent of oligonucleotide hybridization (52). The substitution of adenine by 2-aminopurine in DNA is reported to have only a small duplex-destabilizing effect, measured as 0.5 ± 0.9 kcal/mol at 30°C in 1 M NaCl (53).

Because this work is based on the use of oligonucleotides containing a single 2-aminopurine residue, we proceeded to test the effect of this substitution in the context of RNA sequences at high salt concentration. By measuring melting temperatures of oligonucleotides incorporating 2Ap, we estimated a destabilization effect of ≈ 0.2 kcal/mol at 37°C (see Supplementary Data 1). This effect is therefore negligible at room temperature and is within typical measurement error. In light of this result, we assume that oligonucleotides containing a single 2Ap are a good proxy for the properties of unmodified RNA (54).

### Association kinetics of short oligoribonucleotides

Upon mixing of multiple oligonucleotides in solution they will quickly anneal to form a mixture of all possible double helices resulting from base-pairing. The initial abundance of these duplexes in a competition scenario will reflect the relative hybridization rates (*k*_*on*_) for any pair of strands. It follows that characterization of the behavior of any mixture of short RNA oligonucleotides requires the measurement and modeling of *k*_*on*_, which we address in this first section, where we aim to provide the tools to readily calculate hybridization kinetics for any given pair of sequences of interest and thereby determine the initial configuration of an arbitrary collection of RNA strands.

The sequences used in this work were designed to minimize secondary structures, and map onto six template sequences with CG contents ranging from 0.0 to 0.8 (see Supplementary Data 8 for a list of all sequences). For each template, we measured the kinetics of binding of a series of complementary oligonucleotides of different lengths at concentrations typically ranging from ≈ 0.5 µM to ≈ 5 µM as shown in Figure 2a. The association rates determined through least squares fitting to a second order reaction scheme yielded the values shown in Figure 2b, where a positive dependence of the hybridization rates on oligonucleotide length can be observed.

**Figure 2.**
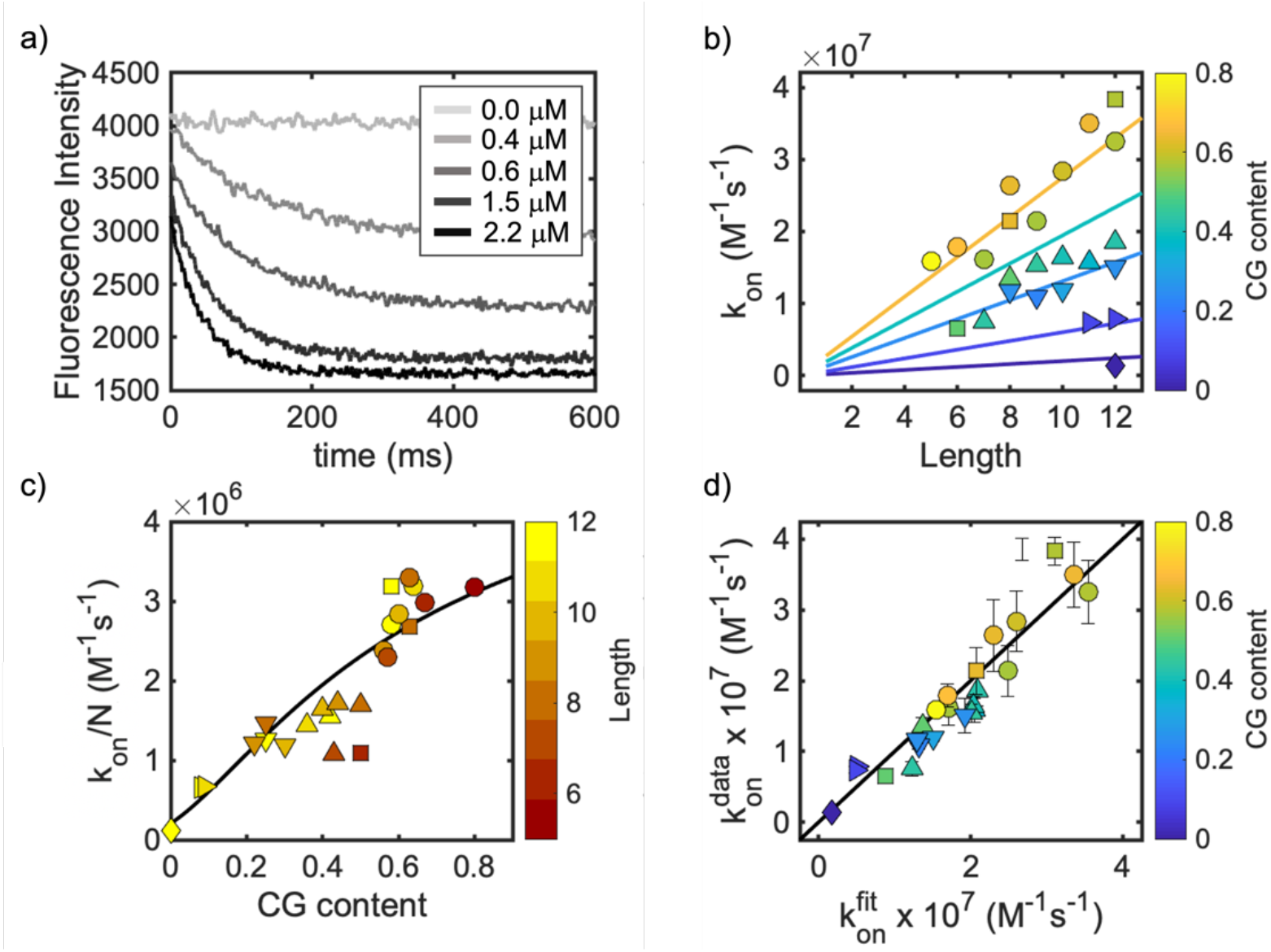
Association kinetics of short RNA oligonucleotides. (a) Example of hybridization of the 8 nt long oligonucleotide K8 to the KC* template sequence (at 0.5 µM) as a function of its concentration. (b) Experimentally derived *k*_*on*_ for short oligonucleotides show a linear dependence on their length. Different symbols are used for experiments performed using different templates as indicated in the legend. Continuous lines represent predicted *k*_*on*_ values computed from Eq. 10 with *K*_*D*_ calculated from Eq. 11. Line colors match colormap on right. (c) length-normalized hybridization rates (M^-1^s^-1^) and approximating function computed form Eq. 10 and Eq. 11. Deviations from the black line are likely due to sequence complexity not captured by an average *K*_*D*_. (d) Correlation plot for calculated *k*_*on*_ using best fit *k*_*bi*_ and *k*_*uni*_ versus experimentally derived *k*_*on*_. *k*_*on*_^*fit*^ is computed using Eq. 9 with *k*_*bi*_ = 1.6 × 10^7^ M^-1^s^-1^ and *k*_*uni*_ = 2.1 × 10^5^ s^-1^. Error bars are shared among data in panels b, c and d and only shown in the latter for clarity.

To model RNA binding kinetics, we used two different approaches: an analytic treatment based on a simplified model and a more exact numerical treatment of the complete process of oligonucleotide hybridization; both approaches were developed based on previously published work in which annealing is considered to begin with the formation of a single base-pair, followed by the subsequent formation (or loss) of additional base-pairs (34,35,55).

In our simplified model, we treat the binding of two oligonucleotides as a multiple-step process: the oligonucleotides first diffuse and collide in the proper orientation to form an initial base-pair with a bimolecular rate *k*_*bi*_. After the initial contact is formed, new base-pairs can be progressively added or removed, with the formation of new base-pairs proceeding at a fixed unimolecular rate *k*_*uni*_. Reverse steps for removal of a base-pair proceed with a rate 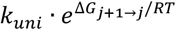, with ΔG being the free energy difference between the state with j+1 bound base-pairs and the adjacent state with j bound base-pairs. Analogously, the rate of detachment of two strands when connected by a single base-pairs is given by 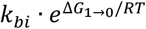, which is equivalent to *k*_*bi*_ · *K*_*D*_, with *K*_*D*_ being the dissociation constant for the two strands linked only by the single base-pair. This whole process can be expressed using the following reaction scheme:

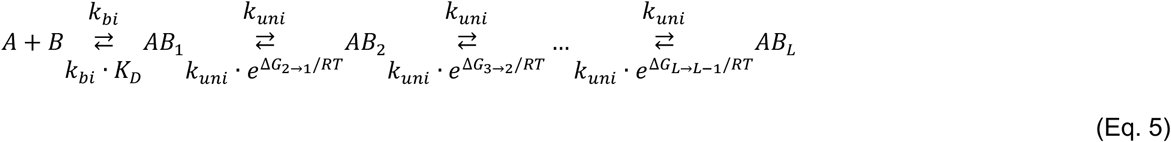

Where AB_j_ represents a bound state between the A and B strands linked by j base-pairs.

From this reaction scheme we can define *k*_*on*_ as the following summation for every i-th initial contact over the total number of initial contacts (N), with N equal to the length of the shortest of the two strands that attempt annealing:

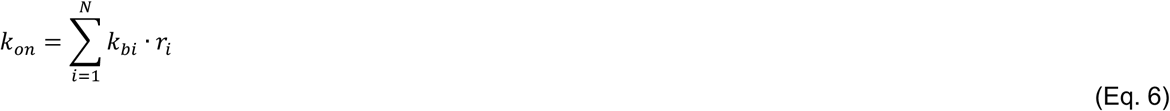

where *r*_*i*_ is the probability of complete zippering after the first i-th contact has been established with a bimolecular rate *k*_*bi*_. Ideally the contact-dependent probability function *r* should take into account every possible trajectory that the two oligonucleotides can follow in order to move from the initial state to the completely annealed state. Because this is impossible to calculate analytically, we approximated it here by considering only the addition and removal of adjacent base-pairs with no bulges or internal loops (34,55). One important consequence of this approximation is that all nucleation events considered must be in-register (i.e. the initial base pairs will also be present in the final duplex), since all off-register events have no way of correcting their trajectory to align as they zipper.

This simplified the problem, allowing us to define this probability function *r* as the product of two terms, *pn* and *pz*. We first defined the probability (*pn*) of going from an AB_1_ to an AB_2_ state from a given i-th initial contact according to the reaction scheme previously shown, and considering that for every initial binding site except the ones occurring at the termini there are two possible zippering directions:

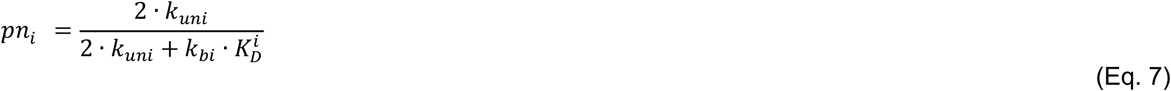

This function determines the probability of reaching the state AB_2_, after which the zippering can proceed as a random walk with probability (*pz*) of undergoing a new step equal to:

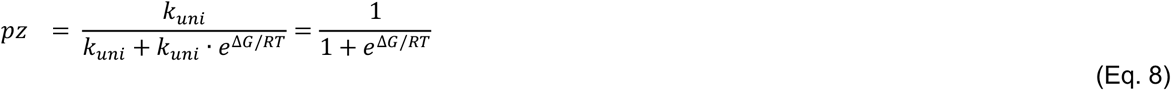

with ΔG calculated as the energy difference between the paired and unpaired states as previously defined. We computed (based on NN thermodynamic parameters) two extreme *pz* values for the probability of propagating the zippering by a single base-pair, finding values approximately equal to 0.84 for f_CG_ = 0 and 0.99 for f_CG_ = 1. Given the high chance of this event, the overall success in hybridizing two strands will be heavily dominated by the probability *pn* of formation of the second base-pair after the initial contact. However, if we assume that the probability of successful zippering is controlled uniquely by *pn*, we will overestimate *r*, since we would neglect the probability of any failing trajectory after the state AB_2_ has been reached.

We calculate this error as follows: we define *pz*^*fail*^ as the overall chance of failing the zippering and eventually falling back to the AB_1_ state after states AB_2_, AB_3_, AB_4_ etc. have been reached. If *pz*^*fail*^ is close to zero, we can effectively neglect *pz* and consider that the overall success rate is controlled uniquely by *pn*. This 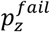 value could be calculated through the Gambler’s Ruin analysis (56), resulting in a probability of ∼16% of falling back to AB_1_ after the second nucleobase is bound for the f_CG_ = 0 case (for f_CG_ = 1 this value is lower than 0.1%), plus an additional smaller probability of ∼3% of falling back to AB_1_ from AB_3_, AB_4_ and so on. It follows that by considering the annealing as successful once AB_2_ is reached, we will incur a maximum error in estimating *r* which is lower than 19% in the worst-case scenario, and specifically equal to *p*^*fail*^ · (1 ∼ *pn*).

Since we have established that the formation of a second adjacent base-pair dominates the overall probability of complete zippering, it follows that the formation of two adjacent base-pairs can be effectively defined as the limiting nucleation event. We therefore approximate *pz* to 1 and redefine *k*_*on*_ using the following equation:

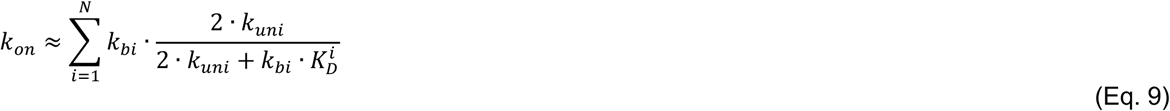

Fitting our experimentally derived k_on_ data with this analytical model allowed us to reproduce measured *k*_*on*_ values with good accuracy (Pearson coefficient = 0.94) and no outliers, with the best fitting *k*_*bi*_ equal to 1.6 × 10^7^ M^-1^s^-1^ and *k*_*uni*_ equal to 2.1 × 10^5^ s^-1^. It is important to note that, while our function is sensitive to the value of *k*_*uni*_, it is not equally constrained by *k*_*bi*_, generating good predictions even when *k*_*bi*_ is much larger than our best fitting value. It follows that even if these parameters can be used to calculate hybridization rates, their physical interpretation should be treated with caution.

Since the computation of the binding energy and *K*_*D*_ for every possible initial contact is complicated by contributions from neighboring but non-base-paired nucleotides, we present here a simple function to approximate it starting from the CG content of the complementary stretch, that can be readily used without additional tools. To do so, we assumed that a single average value *r* describes the probability of successful annealing when any among the possible initial N contacts are formed, where N is the length of the shortest of the two complementary strands having CG content equal to f_CG_. In this case, we can calculate a length-normalized *k*_*on*_ as follows:

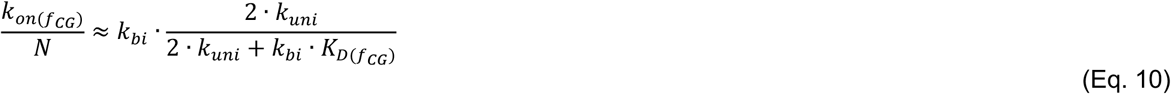

Since there is no straightforward function to compute a priori the free energy of a base-pair, we provide here an empirical function that relates the CG content of the RNA strand to the apparent ΔG of its initial contacts, extracted by expressing Eq. 10 as a function of *K*_*D*_ and fitting our dataset. The following resulting equation can be used to compute *K*_*D*_ in Eq. 10:

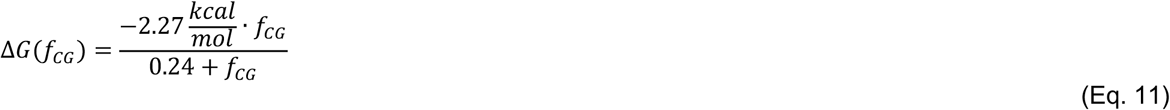

*k*_*on*_ parameters computed using Eq. 10 and 11 are plotted as straight lines in Figure 2b, producing predictions that correlate well with experimentally derived *k*_*on*_ data (Pearson coefficient = 0.91).

With a simplified analytical model in hand, we tested our experimentally derived data against a more complete stochastic model that accounts for bulges and internal loops during the hybridization, which are expected to some degree according to coarse-grained simulations in repetitive sequences (28,36,57). To do so, we implemented in Python the approach established by Schaeffer and released as a software program called Multistrand (35) which models the annealing process as a trajectory over the complete Markov-chain of bound states among two strands, an algorithm based upon the work of Flamm and colleagues for RNA folding (58). Every transition between two states differs by a single nucleotide – either removed or added – and every state is characterized by a defined pattern of bound nucleotides, whose free energy is calculated using NUPACK 4.0, a powerful software package developed by the Pierce group to compute the thermodynamics of systems containing multiple oligonucleotides (59). One advantage of such approach is that it explicitly accounts for every step of the annealing process and should give a reasonable success rate in producing the aligned minimum free-energy structure when starting from an off-register initiation event. Using the publicly available version of Multistrand, we found that even though the overall trend of our data is reproduced (Pearson coefficient = 0.66), there are many poorly predicted data points. In particular, the f_CG_ = 0 sequence has a highly (20 fold) overestimated *k*_*o*n_. Given the low complexity of this sequence and the high number of off-register sites available to initiate its hybridization, the outcome of Multistrand suggests that the success rate for off-register trajectories may be overestimated with the current parametrization.

One important difference between the reaction scheme employed in stochastic simulations on nucleic acids by Schaeffer and Flamm and the simplified analytical model previously established in Eq. 5 is the implementation of the Metropolis rate method for zippering. Briefly, in the stochastic simulation the rate for any transition between two states that does not result in the detachment of the two strands is equal to *k*_*un*i_ for ΔG < 0 (favorable transitions) or *k*_*uni*_ * e^(-G/RT)^ for ΔG > 0 (unfavorable transitions). To obtain hybridization rates through stochastic simulations, we implemented the “first-step” approach established by Schaeffer (35): the two oligonucleotides were initialized in a complex connected by a single base-pair picked randomly among all initial in-register and off-register nucleotides. The system then evolves using the Gillespie algorithm (60), picking a new state among all possible states that differ by a single base-pair with a probability scaling on their rates. This process was iterated to produce a trajectory with the code stopping if the complex successfully reaches its minimum free-energy structure, or failing when the two strands detach.

By resampling hybridization trajectories many times, we built up the success probability *r* for the annealing of two strands. Since the trajectories were solved using the Gillespie algorithm, we could compute an accurate *k*_*on*_ that accounts for the time spent by the two strands in solution before colliding and the time spent in each zippering process (35), leading either to success with an average time spent *τ*_*succ*_ or to failure with an average time spent *τ*_*fail*_.

We should define first a collision time, which is the time spent by two oligonucleotides in solution before forming one among N possible initial contacts:

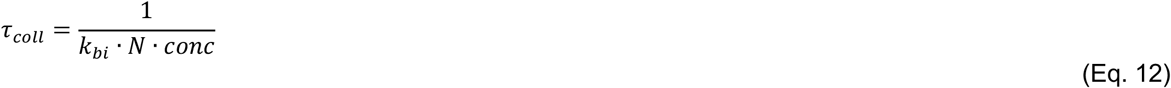

From this we could compute the *k*_*on*_ taking into account the time spent by the strands to form the initial base-pair and the time spent in failed attempts before their successful annealing:

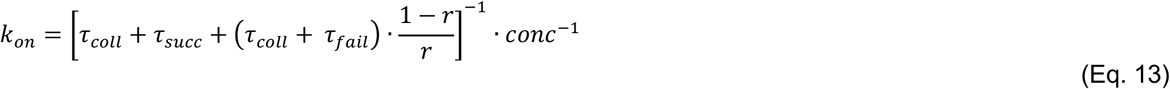

where *conc* is the oligonucleotide concentration (in our case we use 1 *µM*), N the total number of initial binding sites and *r* the average success rate over all initial interactions. By doing a grid search, we found the best fitting *k*_*bi*_ equal to 3.5 × 10^6^ M^-1^s^-1^ and *k*_*uni*_ equal to 4.25 × 10^5^ s^-1^ (Pearson coefficient = 0.91), producing the simulated *k*_*on*_ values plotted against our experimental data in Figure 3a. Among the advantages of solving the complete Markov-chain for oligonucleotide annealing is that we could look in more detail at the role of in-register and off-register binding events(28,57) and their trajectories. Here we discuss annealing pathways considered as the set of unique binding configurations encountered during a trajectory, each one with a certain free-energy difference calculated from the reference unbound state and a minimal formation time, defined as the time elapsed from the initial contact until the given state is first reached. Looking at an in-register pathway (Figure 3b) we can see that it is a smooth function, with the annealing process resolving within a few microseconds. In contrast, with off-register contacts, we generally predicted a low success rate in hybridizing if the sequences have a low CG content (< 0.5) or they are relatively short (< 8nt). For longer sequences with high CG content (>= 0.5), we have found a non-negligible probability of producing the minimum free-energy structure starting from an off-register site - as long as the offset is not too extreme - with zippering passing through a more complex pathway with kinetic traps visible as in the case depicted in Figure 3c, where a pathway leading to the correct annealing of two strands starting from an off-register nucleation is plotted as an energy-time function.

**Figure 3.**
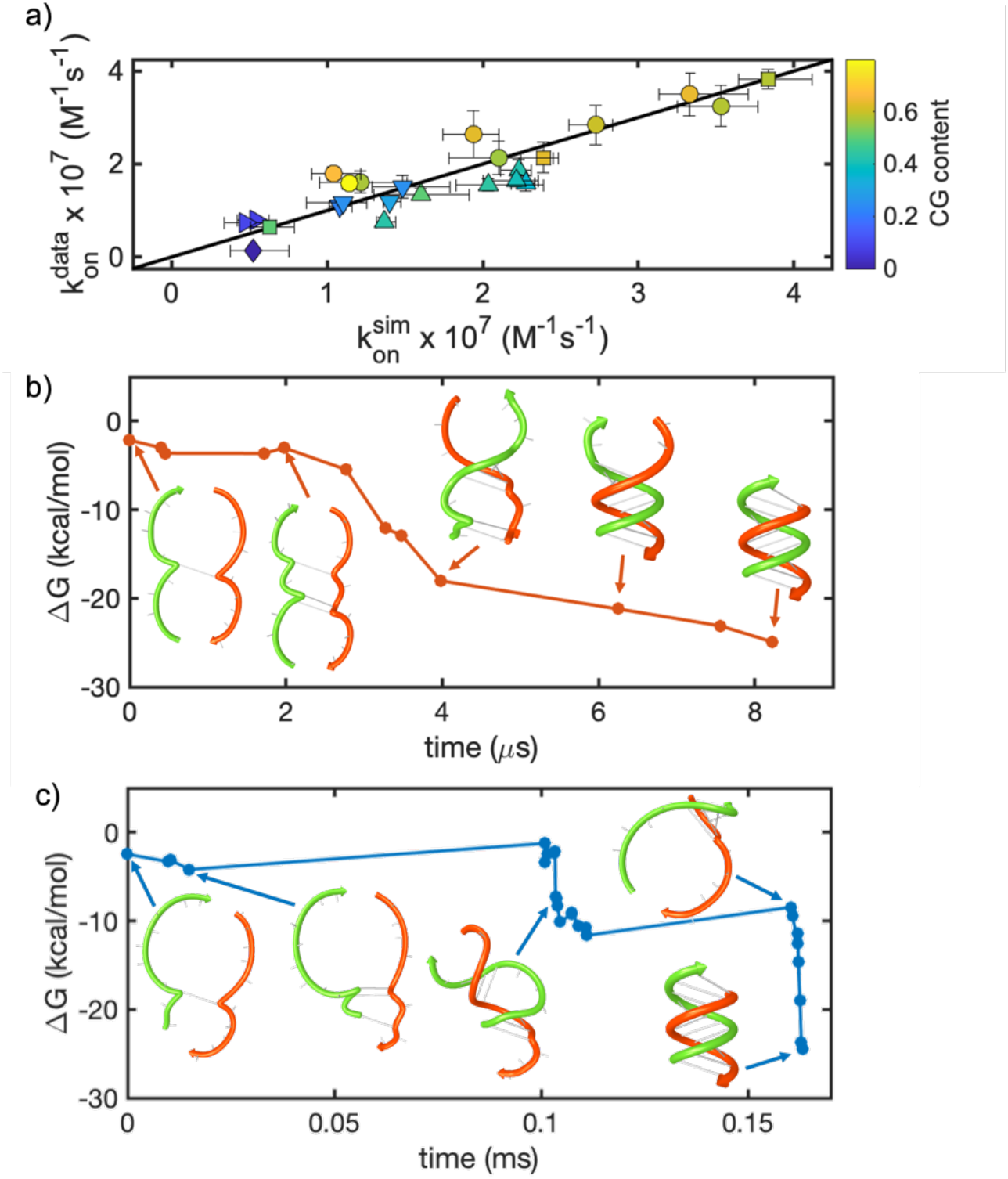
Stochastic simulations for the association kinetics of short RNA oligonucleotides. (a) Correlation plot for calculated *k*_*on*_ versus experimentally determined *k*_*on*_. Calculated values were produced using *k*_*bi*_ = 3.5 × 10^6^ M^-1^s^-1^ and *k*_*uni*_ = 4.25 × 10^5^ s^-1^, sampling each initial binding site ten times and averaging over three replicates. Error bars on the x-axis represent standard deviations between the three replicates. Symbols used here refer to the same templates as the symbols used in Figure 2. (b, c) Annealing pathways for in-register (b) and off-register (c) contacts. In the first case we see a smooth pathway to the energy minimum, for the latter case the pathway shows kinetic traps solved through the correct realignment of the two strands. As soon as the two strands become correctly aligned, they zipper in a few microseconds. Selected pairing configurations along the two pathways are sketched using NUPACK web-based utilities to give a clearer representation of the different stages leading to a successful annealing.

In summary, our analysis of the annealing process of two short RNA oligonucleotides has allowed us to construct a simple model that returns accurate values of *k*_*on*_ using just two parameters (length and CG fraction), constituting the first step in producing a practical predictive model for the evolution over time of more complex systems of short RNA oligonucleotides.

### Dissociation kinetics of short oligoribonucleotides

After the initial configuration of double helices in a mixture of short RNA oligonucleotides is established according to the hybridization rates for all pairs of strands, the duplexes will eventually separate after a certain amount of time governed by their *k*_*off*_. This rate is crucial in determining the relaxation time-scale for any complex mixture and it is explored in this section.

Having measured *k*_*on*_ for our set of oligonucleotides and shown that these measured rates are in good agreement with a simple predictive model, we proceeded to determine the corresponding *k*_*off*_ rates from their dissociation constants as calculated from the NN model for oligonucleotides. This approach is reliable as long as duplex formation can be approximated as a two-states process, as already validated in the field of DNA hybridization (23,27). Before proceeding, we checked the accuracy of NN predictions of *K*_*D*_ for RNA duplexes at room temperature, since systematic discrepancies on the order of ∼1 kcal/mol are often found in studies on DNA (23,25,27). We obtained NN-predictions of the ensemble free energy (which includes the effects of frayed termini and other suboptimal configurations) by performing simulated melting experiments with NUPACK. We compared these values with our experimental data obtained by measuring the free energy of binding in a subset of our sequences through (i) direct *K*_*D*_ measurement through static binding experiments at room temperature *via* fluorescence spectroscopy, (ii) UV melting experiments, (iii) fluorescence melting experiments and (iv) kinetic binding measurements, finding that our measured ensemble free energy agrees very well with the predictions of the NN model (Pearson coefficient = 0.98, see Supplementary Data 3).

Since NN-derived ΔGs are therefore reliable for the sequences used in this work, we calculated the *k*_*off*_ for all of our studied oligonucleotides as *k*_*on*_⋅*K*_*D*_, where *K*_*D*_ was computed as e^ΔG/(R⋅T)^ and ΔG is the free energy difference between the unbound and bound state. Data obtained in this way are presented as a function of oligonucleotide length (Figure 4a) and of the free energy of binding as a comparison with *k*_*on*_ (Figure 4b and 4c).

**Figure 4.**
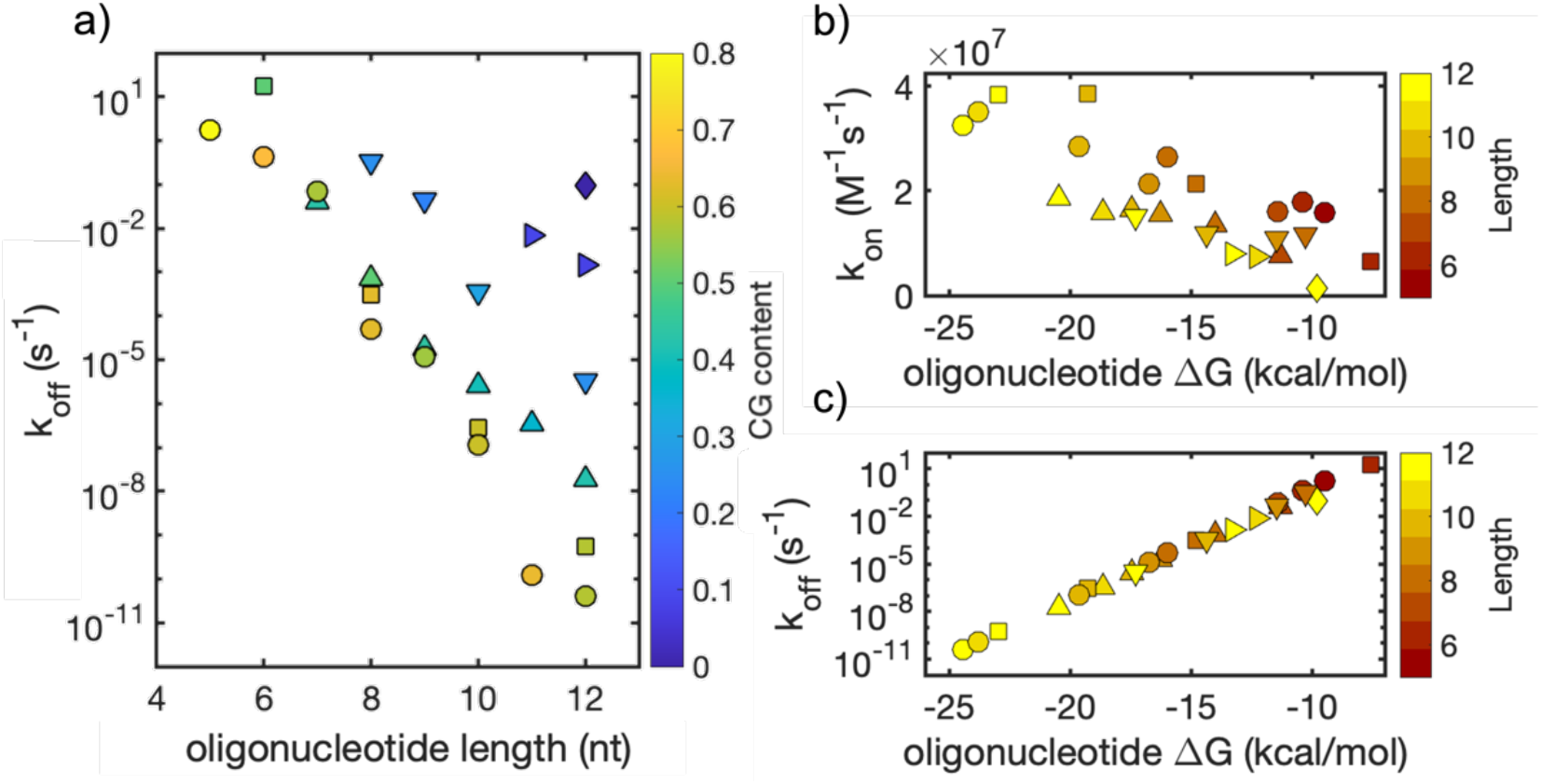
Kinetics of dissociation and association of short RNA oligonucleotides. *k*_*on*_ was measured as previously described in this work, while *k*_*off*_ was calculated as *k*_*on*_⋅*K*_*D*_. (a) *k*_*off*_ for the oligonucleotides studied in this work as a function of length shows a poor correlation and emphasizes the dramatic scaling and role of CG content. (b) *k*_*on*_ as a function of ΔG shows a linear dependence. (c) k_off_ as a function of ΔG shows a strong exponential correlation.

As expected, we observe a strong exponential correlation of *k*_*off*_ with ΔG, in agreement with the literature and the linear relationship between the activation energy for dissociation and ΔG (25,29). An equally clear linear correlation of *k*_*on*_ with ΔG can be observed, which is a direct consequence of the relationship stated in Eq. 9. It follows that the main contributor to the overall duplex stability of the RNA oligonucleotide is *k*_*off*_, whose values vary by roughly ten orders of magnitude for the oligonucleotides studied in this work. It is important to note that *k*_*off*_ in RNA is much lower than that for DNA (29,39), with the parameter quickly reaching timescales much longer than common processes of interest in prebiotic and biological contexts. A consequence of this phenomenon is that equilibration processes for a solution of many oligonucleotides might take several years unless mechanisms to speed up this process intervene, such as strand displacement.

### Toehold mediated strand displacement

Whenever a double helix and a single stranded oligonucleotide are present in solution, the equilibration will not be necessarily limited by the *k*_*off*_ of the duplex and could be greatly sped up due to strand displacement, a process that will inevitably happen in a competing scenario.

Strand displacement is a multi-step process where an oligonucleotide (A) bound to a template (B) is replaced by a third oligonucleotide (C). During the strand displacement process, the incoming displacing oligonucleotide binds to an available complementary stretch on the template, the so-called toehold, and undergoes branch migration, which leads to the release of the previously bound strand. We designed our sequences so that the displaced oligonucleotide is unlikely to rebind to the template since the latter is missing a useful toehold once bound to C. A complete reaction scheme for a system undergoing strand displacement should take into account association and dissociation of every species as depicted in Figure 5a.

**Figure 5.**
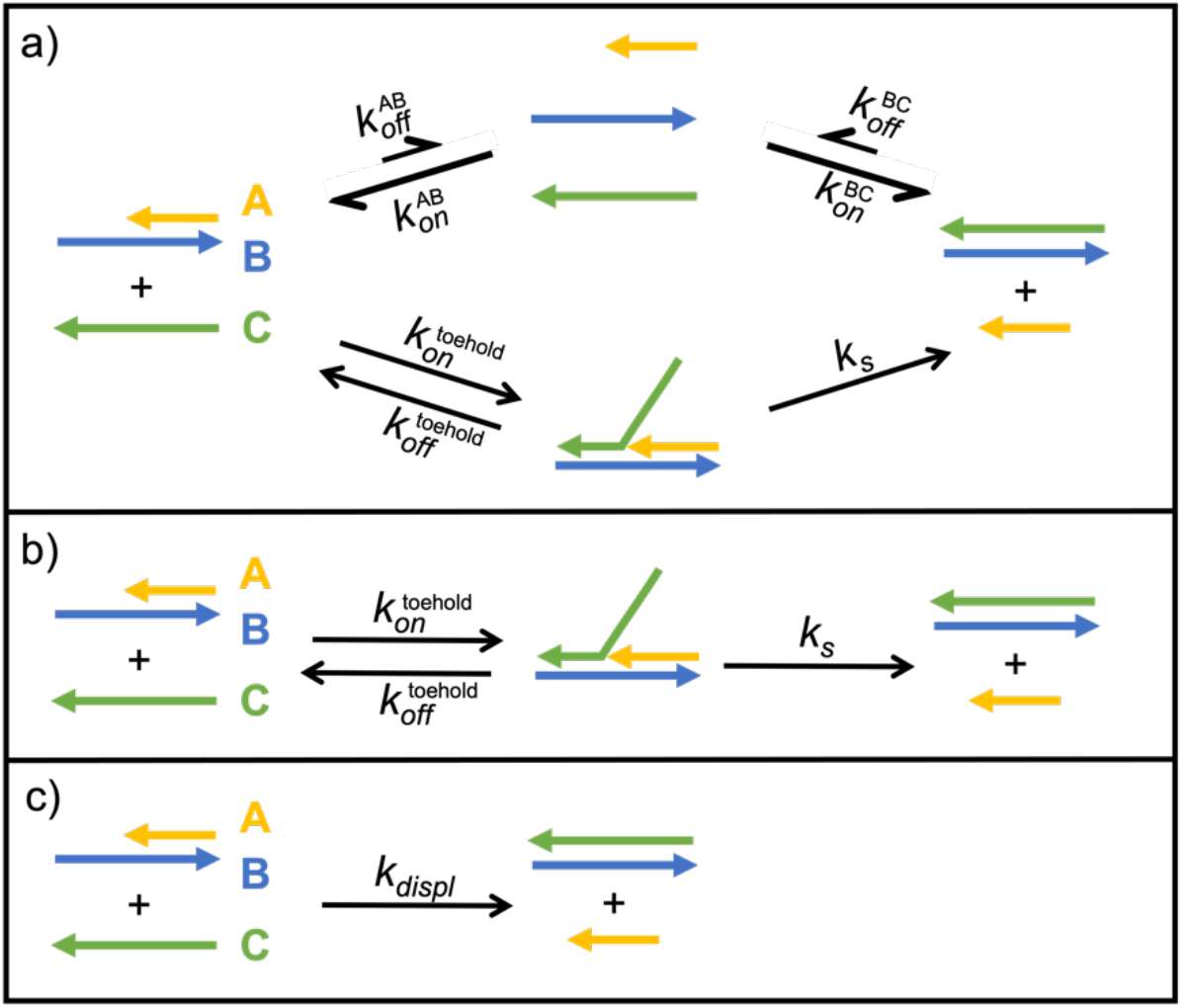
Reaction scheme for toehold-mediated strand displacement. (a) Complete reaction scheme. (b) Reaction scheme under the condition of negligible dissociation of duplexes. (c) Simplified reaction scheme valid when a rapid pre-equilibrium is established between the displacing oligonucleotide and the template.

If the strand displacement occurs on a timescale much shorter than the lifetime (τ) of the AB duplex, we can neglect the dissociation of both AB and BC (*τ*_BC_ > *τ*_AB_ by design) so the reaction scheme can be reduced to a two-step reaction as shown in Figure 5b. Moreover, if the toehold binding equilibrates on a much faster time scale than the strand displacement reaction, which is reasonable for the short toehold lengths explored in this work, we can approximate the strand displacement process as a much simpler second order reaction, as shown in Figure 5c.

This simple model treats strand displacement as a bimolecular reaction with second order rate *k*_*displ*_ equal to *k*_*s*_*⋅K*_*A*_, where *K*_*A*_ is the association constant of the toehold (*K*_*A*_ = [ABC][AB]^-1^[C]^-1^). In our experiments we monitored the evolution of the system by following the 2Ap fluorescence signal. Because [ABC] << [C] using short toeholds, even if 2Ap is placed in the toehold domain, the quenching measured in our experiment derives solely from formation of the BC duplex product. For the very same reason, because [ABC] is too low to be determined experimentally, we cannot directly measure *K*_*A*_. To overcome this limitation we take advantage of NUPACK following the approach of Zhang & Winfree (44). Using NUPACK we can compute the free energy of user-defined structures and determine the difference in energy from states “AB + C” to state “ABC”. These values can be used to calculate the toehold binding energy (ΔG_toehold_) as ΔG_AB+C_ – ΔG_ABC_ and the association constant for the toehold binding (*K*_*A*_) equal to e^(ΔGtoehold/RT)^.

For the case of strand displacement without an available toehold, we assume that strand displacement proceeds through the binding of C to the terminal nucleotides of the AB duplex, which are partially available due to the fraying of the duplex ends (61). In this case the strand displacement rate will be the sum of the displacement rates at the two termini, k_s_⋅(K_A_^1^ + K_A_^2^), with an apparent binding energy of the invading strand to the toehold equal to:

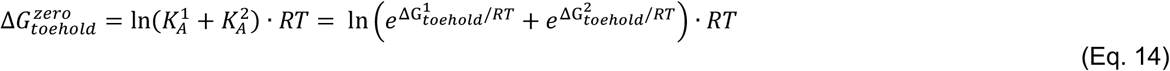

According to the model used here, a plot of *k*_*displ*_ versus ΔG_toehold_ should follow the exponential function *k*_*s*_⋅e^(-ΔGtoehold/RT)^ that we used to fit our dataset obtained from a series of experiments performed using toeholds of lengths ranging from 0 nt to 5 nt and positioned either on 5′ or 3′ terminus of the template.

As expected, we find that longer toeholds induce faster strand displacement up to saturating values of ≈ 2⋅10^6^ M^-1^s^-1^, while for toehold binding energies lower than 7kcal/mol, our data yields a best fit of *k*_*s*_ = 12 s^-1^ (Figure 6b) with no significant difference between toeholds positioned on the 5′ or 3′ terminus of the template.

**Figure 6.**
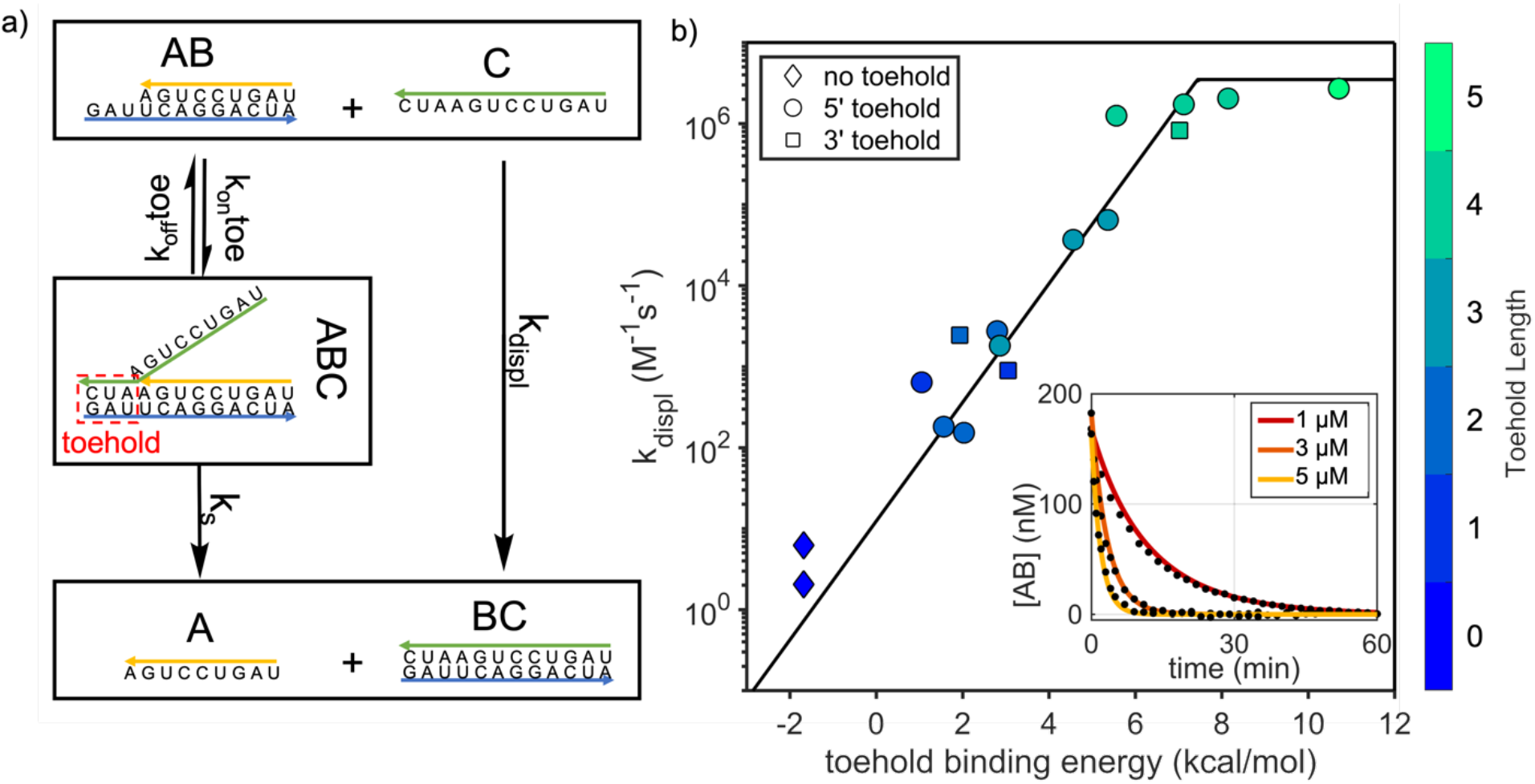
Strand displacement in short RNA oligonucleotides. (a) Reaction scheme for strand displacement. Under our conditions the two-step reaction can be simplified as a single bimolecular reaction with rate *k*_*displ*_. (b) *k*_*displ*_ for short oligonucleotides has an exponential dependence on toehold binding affinity. Toehold binding affinity for the case with no toehold is derived as described in main text. The best fit for *k*_*s*_ = 12 s^-1^ is shown (black line). Saturation value for plotting purposes has been set at *k*_*bi*_ (3.5⋅10^6^ M^-1^s^-1^). Error bars are smaller than symbols size and are thus not shown. Inset depicts strand displacement time traces for different [C] for a 3nt long toehold (ΔG_toehold_ = 2.86 kcal/mol). Dots are experimental data and continuous lines are the result of a global fit for *k*_*displ*_ = 1.81⋅10^3^ M^-1^s^-1^.

### Approach to equilibrium of mixtures of oligonucleotides

Having measured the kinetic parameters governing the time-evolution of sets of oligonucleotides, we asked whether our approach to modeling the kinetics of annealing of short oligonucleotides could be used to predict the behavior of more complex mixtures. For each experiment we mixed four oligonucleotides: two short oligonucleotides (A_S_, B_S_) and two long oligonucleotides (A_L_, B_L_) designed so that all A-type and B-type molecules could bind to each other (see Figure 1 for sketch of the experimental design). We followed the competition between different binding interactions by monitoring the disappearance of the A_S_B_L_ (and appearance of A_L_B_L_) duplex by placing 2Ap in a region of B_L_ where it is selectively quenched by A_L_; the duplex of B_L_ with A_S_ leaves 2Ap in an overhang and therefore unpaired and unquenched. Once again, we exploited the properties of 2Ap to track the evolution over time of competing oligonucleotides. After mixing the single stranded oligonucleotides at a concentration on the order of 1 µM, the fluorescence signal rapidly drops due to the immediate formation of all possible duplexes: A_S_B_S_, A_L_B_L_, A_L_B_S_ and A_S_B_L_ coexisting in solution at different concentrations. This first process is too fast to be captured by hand mixing and thus is not shown in our time traces. After this initial rapid hybridization, the system is still out of equilibrium and evolves over time until it converges to its energy minimum, which corresponds to a state where all the duplexes in solution are A_S_B_S_ and A_L_B_L_, while less stable duplexes A_L_B_S_ and A_S_B_L_ are absent. As the experiment progressed, we observed more and more quenching of the fluorescence emission of B_L_ due to its annealing to A_L_. At all times after the initial rapid annealing, it must be true that [A_S_B_S_] ≈ [A_L_B_L_] and [A_S_B_L_] ≈ [A_L_B_S_]. We could therefore determine the time course of each component of the system in the same experiment. In a typical measurement, we see the fluorescence dropping on a time scale that varies drastically depending on the oligonucleotide sequences and concentrations, with the presence of longer toeholds for strand displacement speeding up the equilibration in a way that can be tuned on time scales varying by orders of magnitude.

After the duplexes are quickly formed in solution with a relative abundance depending on their hybridization rate, different pathways are expected to drive the quenching of the fluorescent signal and equilibration of the system: (i) direct hybridization of A_L_ and B_L_ oligonucleotides as they become briefly available after detaching from their B_S_ and A_S_ counterparts (Figure 7a), a process in direct competition with

**Figure 7.**
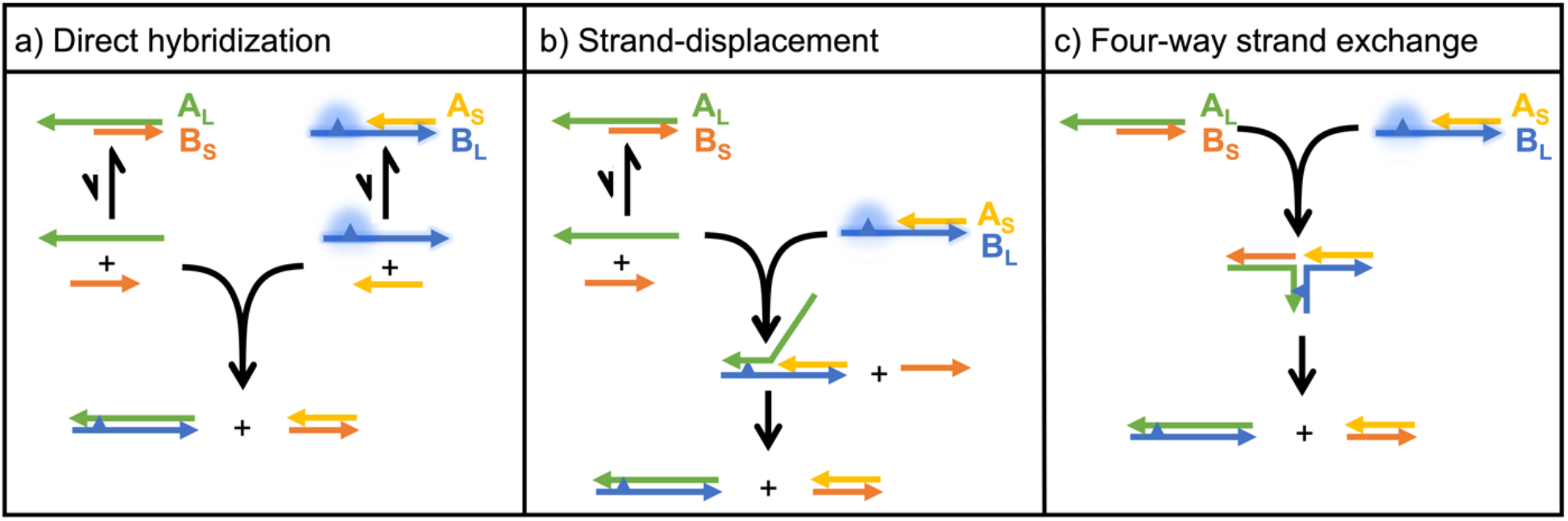
Equilibration pathways expected to determine the evolution of A_S_B_L_ in complex mixtures of competing oligonucleotides.

(ii) strand displacement from available A_L_ in solution due to an imbalance of the initial concentration of the different species or because they have become available after briefly detaching from their complementary B_S_ strand (Figure 7b) and (iii) four-way strand exchange, driving the direct substitution of two strands mediated by the interaction of the overhangs of two A_S_B_L_ and A_L_B_S_ duplexes, without the need for prior dissociation (Figure 7c).

According to the study by Nadine (62) on DNA four-way branch migration, we expected the latter phenomenon to be extremely slow. To test whether this pathway was relevant in our case studies, we prepared mixtures of pre-annealed A_S_B_L_ and A_L_B_S_ at different total concentrations ([A_S_B_L_] = [A_L_B_S_]) and using sequences that would expose 4nt long complementary overhangs, the longest studied here. If a direct reaction pathway such as the four-way branch exchange occurs, the equilibration of the system should exhibit a concentration-dependent rate. However, our measurements clearly show a concentration-independence of the equilibration rates, allowing us to rule out the four-way branch exchange pathway (see Supplementary Data 6). Our data set an upper limit for the rate of the four-way strand exchange process, which must be slower than ≈ 10^2^ M^-1^s^-1^ for duplexes with 4nt long overhangs.

Having ruled out the third equilibration pathway, we focused on the first two, that should be well described by the parameters characterized in this work. To model our time course experiments, we first calculated *k*_*on*_, *k*_*displ*_ and *k*_*off*_ as described in the previous sections, and then numerically solved a set of differential equations describing the system using the calculated parameters, varying them within their experimental error range (see Supplementary Data 7 for the complete system of differential equations). In Figure 8, we show experimental results compared with our predictions depicted as shaded areas. The time courses numerically computed using parameters calculated as described in this work do approximate the evolution of the different species, capturing equilibration times ranging from a few minutes up to several hours. For each case study, we determined the relative contribution of pathways (i) and (ii) toward equilibration after the first quick hybridization has been established, computed as the relative amount of A_L_B_L_ produced through the two pathways and expressed as percentages in the right panel of Figure 8. We can describe two general cases for the equilibration of these mixtures: (i) for unbalanced mixtures having concentration of A_L_ larger than A_S_, B_S_ and B_L_ (Figure 8b, c) we can in principle push the equilibration rate up to the toehold binding rate by increasing the toehold length and A_L_ concentration; for balanced mixtures (Figure 8a, d) the relaxation timescale shows a lower limit corresponding to the lifetime of A_S_ and B_S_-containing duplexes. The equilibration in such case can effectively coincide with the lifetime of the A_S_B_L_ duplex (≈ 1 hour) when strand displacement is very fast (Figure 8a) or it can be dominated by the direct hybridization of transiently freed A_L_ and B_L_ when strand displacement is slow (Figure 8d), thus extending the equilibration timescale up to several duplex lifetimes.

**Figure 8.**
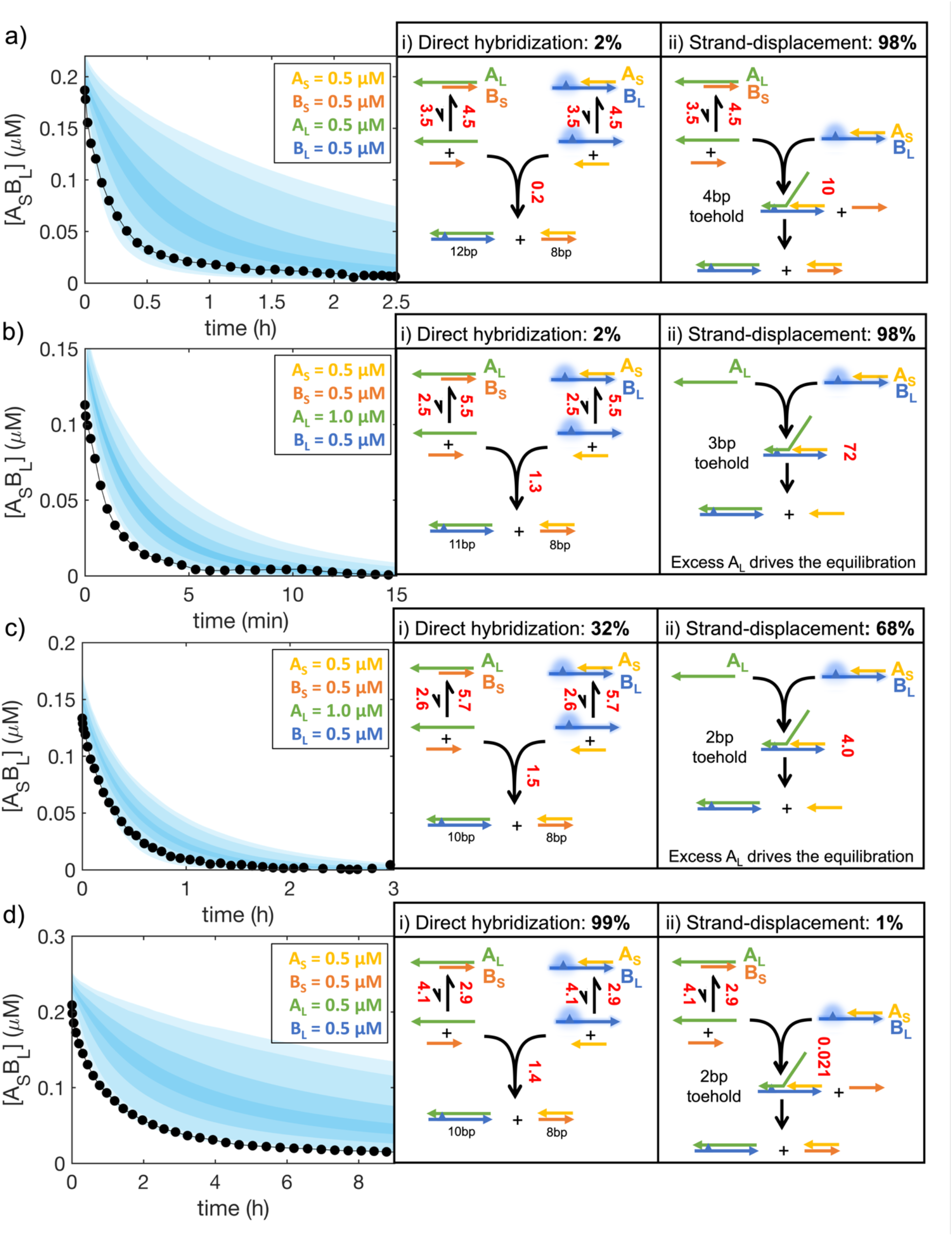
Examples of time traces for the evolution of A_S_B_L_ in complex mixtures of competing oligonucleotides (black) compared with predictions from our modeling (shaded area). The measured signal is coming from the species A_S_B_L_ and disappears when the solution is composed only by A_S_B_S_ and A_L_B_L_ following the pathways depicted on the right. For each pathway the relative contribution to equilibration is given as a percentage of A_L_B_L_ produced after the initial rapid hybridization. Concentrations of sequences used are specified for each panel. Numbers in red next to each reaction arrow specify the corresponding instantaneous rates in units of nM/min, calculated right after the initial duplexes are established through the rapid hybridization of the free single stranded RNA (typically a few seconds after mixing). Shaded areas enclose trajectories calculated by resampling the kinetic parameters within their estimated confidence intervals and are drawn to envelope density probabilities equal to 95%, 90%, 60% and 20% of overlapping the real trajectory. Sequences used are specified in Supplementary Data 7.

**Table 1.** Table of names.

| Name | Definition |
| --- | --- |
| $k_{on}$ | Hybridization rate of two oligonucleotides |
| $k_{off}$ | Detaching rate of two oligonucleotides |
| $K_D$ | Dissociation constant |
| $K_A$ | Association constant |
| $k_{bi}$ | Bimolecular collision rate for the formation of a contact between two oligonucleotides |
| $k_{uni}$ | Unimolecular rate for the formation of a new base-pair in a zippering process |
| $r$ | Probability of successfully zippering after a first contact is made between two oligonucleotides |
| $pn$ | Probability of forming a second base-pair adjacent to the first base-pair formed upon contact between two oligonucleotides. |
| $pz$ | Probability of extending an annealed stretch by a single nucleotide after a two base-pair stretch is formed. |
| $\tau_{coll}$ | Average time required for two oligonucleotides to form a first contact. |
| $\tau_{succ}$ | Average time required for two oligonucleotides to undergo a successful zippering trajectory after the first contact is established. |
| $\tau_{fail}$ | Average time required for two oligonucleotides to undergo a failing zippering trajectory after the first contact is established. |
| $k_{displ}$ | Bimolecular rate of toehold mediated strand displacement. |
| $k_s$ | Unimolecular rate of strand displacement after the toehold is bound. |
| $k_{mig}$ | Unimolecular rate for the single step of branch migration during a strand displacement. |

## DISCUSSION

We have examined the kinetics of strand association and dissociation for RNA oligonucleotides having CG content ranging from 0.0 to 0.8 and length between 5 nt and 12 nt. We have provided a simplified analytical model to predict RNA oligonucleotide binding kinetics and validated a Markov-chain model for the description of complete binding trajectories. Our analysis shows that RNA annealing is generally limited by the formation of just two in-register base-pairs in the context of a nucleation-zipper model. Following this result, two base-pairs constitute a nucleation site with a high probability of progressing to complete annealing *via* a fast zippering process. For relatively long oligonucleotides with a high CG content, stochastic simulations show that off-register trajectories become available, with a zippering timescale dominated by the realignment of the two strands. We find that our model is consistent with a single collision rate, supporting the hypothesis of Wetmur and Davidson (34) that any specific nucleation event will occur at a rate that is independent of the length of the oligonucleotide for short oligonucleotides (< 100nt). Since the number of possible in-register nucleation sites is proportional to length, it follows that *k*_*on*_ will show a linear dependence on oligonucleotide length, as observed in this work, and an expected superlinear dependence on length for oligonucleotides having high CG content.

Interestingly, although a sharp drop in hybridization rate for oligonucleotides with fewer than 7 consecutive complementary nucleotides has been observed for both RNA and DNA, leading to the formulation of the so-called “rule of seven” (1), we do not observe a discontinuity in on-rates vs. length at a length of 7, suggesting that a more complicated mismatch-dependent phenomenon might be at play in the work by Cisse and colleagues.

Our model is in agreement with the results of Rejali et al. (25) who showed that a NN-based prediction of *k*_*on*_ is unsuccessful. With the aim of providing a predictive algorithm for the hybridization kinetics of short RNA oligonucleotides, we have shown that it is possible to calculate reliable *k*_*on*_ values using Eq. 10 and 11 as a function of length (N) and CG content, two parameters that are easily determined. With these *k*_*on*_ parameters in hand, we could compute the dissociation rate or *k*_*off*_ of short duplexes from the free energy of binding, calculated using the nearest-neighbor model for RNA. This step is critically dependent on the accuracy of the calculated ΔG, and we found that NUPACK predicted the experimentally measured values with reasonable accuracy. We find a clear dependence of *k*_*off*_ on oligonucleotide length and ΔG in agreement with existing literature (23,25,27,39). As expected *k*_*off*_ for progressively longer RNA duplexes quickly reaches extremely long timescales, making equilibration of complex mixtures unlikely without thermal annealing or some other non-equilibrium relaxation phenomena.

To begin to explore the relaxation time-scales of more complex mixtures of oligoribonucleotides, we first explored strand displacement in RNA duplexes. Our experimental results are in agreement with the coarse-grained simulations of Šulc et al. (45) As previously shown for DNA, we also find that this process is controlled by the binding energy of the invading strand for the template toehold and we have determined a strand displacement rate *k*_*displ*_, equal to *k*_*s*_*⋅K*_*A*_, with a best fit of *k*_*s*_ = 12 s^-1^ for toeholds having ΔG < 7 kcal/mol. Values of the same parameter measured for DNA range from 0.7 s^-1^ to 7 s^-1^(44). Compared to strand displacement in DNA, where the rate plateaus at values close to *k*_*on*_ for a toehold length of ≈ 6 nt, the RNA strand displacement rate in our study approaches limiting values of ≈ 2⋅10^6^ M^-1^s^-1^ using a toehold of 4 nt (when the CG content of the toehold is ≈ 0.5) due to its stronger base pairing. This observation implies that equilibration processes in complex RNA systems will be generally faster than analogous DNA systems if they are dominated by strand displacement. For a more detailed analysis of strand displacement, branch migration can be modeled as a process with a unimolecular rate *k*_*s*_ that can be decomposed using the Gambler’s Ruin analysis as equal to *k*_*mig*_⋅N^-2^ (56), where *k*_*mig*_ is the first-order rate for one elementary branch migration step and N is the length of the duplex region being displaced by branch migration. According to Zhang et al. (44) *k*_*mig*_ in DNA at 1 M NaCl is ≈ 400 s^-1^. For our data, we obtain *k*_*mig*_ equal to ≈ 565 s^-1^, confirming that branch migration occurs at a similar rate for RNA.

Altogether, in this work we have provided a framework for the prediction of the hybridization kinetics of short oligoribonucleotides, and we have addressed the challenge of predicting the time-course evolution of mixtures of competing oligoribonucleotides. We have shown that our approach can be used to compute the evolution of equilibrating solutions containing multiple RNA sequences, which is ultimately dependent on two competing pathways of direct hybridization and strand displacement. While we could rule out the role of four-way strand exchange in our case studies, we can speculate here whether this pathway is going to be relevant when dealing with longer overhangs, that would result in an increase of the four-way strand exchange rate. If we keep the 12nt-long template fixed, we must reduce the duplexed stretch to elongate the overhang, and this will cause a drastic speed up of its *k*_*off*_ and reduction of its lifetime. Using as a reference the branch migration rates published by Nadine (62) and assuming the ones for RNA are comparable (as expected following our results for RNA strand displacement), it is reasonable to expect the equilibration of two duplexes with a long overhang to be ultimately dominated by the first two pathways previously described, so that four-way branch migration is effectively always negligible in mixtures of short RNA oligonucleotides.

By solving a simple system of differential equations, we could predict the lifetime and concentration of every component of the complex mixture, finding that the time required to anneal two complementary strands can be greatly extended by increasing the complexity of the system. While two oligonucleotides typically anneal in a fraction of a second, by varying the composition of a mixture of only four oligonucleotides we could easily manipulate the equilibration time from minutes up to several hours by creating kinetic traps. One important implication of this result is that complexes that are not expected to exist at equilibrium will be present and long-lived when the system initializes in an out-of-equilibrium state. Surprisingly, we find that such duplexes can exist for timescales much longer than their *k*_*off*_-determined lifetimes when their equilibration process is dominated by the direct hybridization pathway. Following this observation, we predict that a system composed by a large collection of duplexes exposing only short toeholds will undergo an exceptionally long equilibration process. This finding is particularly relevant in the context of models for the nonenzymatic replication of genetic information, which is believed to be driven by chemical reactions occurring in highly heterogeneous and complex mixtures of oligoribonucleotides (11,63).

Our work provides the basis for future studies on larger systems of competing strands, and to the design of systems with tunable equilibration rates for desired species. We suggest that a simple extension of our parameters to account for the effect of coaxial stacking on the kinetics of annealing, dissociation and strand displacement will allow this approach to be extended to the study of complexes composed of many strands in large interacting networks of oligonucleotides.

## DATA AVAILABILITY

Oligonucleotide sequences and mixtures studied here are listed in the online supplemental data file accompanying this article.

## SUPPLEMENTARY DATA

Supplementary Data are available at NAR online.

## ACKNOWLEDGEMENTS

We would like to thank Dian Ding, Shriyaa Mittal and Lijun Zhou for valuable discussions and for sharing their virtual circular genome sequence design. We thank Harry Aitken, Brennan Ashwood, Marco Buscaglia, Ben Colville, Daniel Duzdevich, Victor Lelyveld, Kyle Strom, Andrei Tokmakoff and Longfei Wu for comments on the manuscript. M.T. would like to thank Giuliano Zanchetta for introducing him to nucleic acid kinetics.

