## Supplementary Data for "Hybridization kinetics of out-of-equilibrium mixtures of short RNA oligonucleotides"

### 1. 2-AMINOPURINE EFFECT ON RNA THERMODYNAMICS

In this work we used 2-aminopurine as a probe for paired/unpaired state on RNA oligonucleotides. One of the main concerns regarding this approach has been whether 2-aminopurine could have heavily altered the behavior in such a way that it could have not been extended to adenine.

In order to test this hypothesis, we have performed UV melting experiments of a short 5nt oligonucleotide (GCGUG) paired on a template with either adenine or 2-aminopurine in the second position. The results obtained from Van't Hoff analysis are as follows:

| Species: | $\Delta H$ | $\Delta S$ | $\Delta G_{25°C}$ | Buffer |
|---|---|---|---|---|
| 2Ap | -58.00 | 0.1592 | -10.48 | Tris-HCl 200mM, 100mM $MgCl_2$, pH 8 |
| Adenine | -61.88 | 0.1724 | -10.52 | Tris-HCl 200mM, 100mM $MgCl_2$, pH 8 |
| Adenine | -60.20 | 0.1664 | -10.58 | Tris-HCl 5mM, 1M NaCl, pH 7 |

In agreement with literature data on DNA(1) we find a modest destabilizing effect of 2-aminopurine at 37°C, which becomes negligible approaching room temperature. This observation reinforces the validity of our approach and of our results. Moreover, no significant difference could be measured by changing the buffer. Data used for the analysis are shown below:

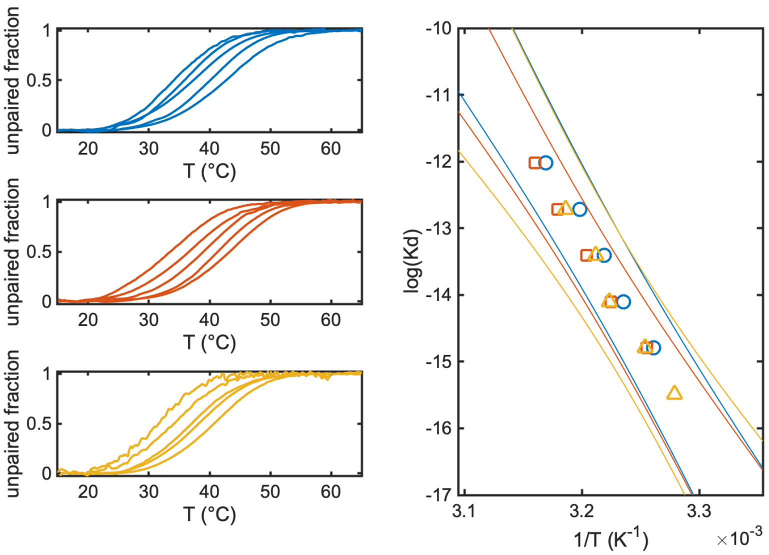

**Supplementary Figure 1.** Top left panel: UV melting curves for 2-aminopurine containing oligo in 200mM Tris-HCl 100mM $MgCl_2$ pH 8. Central left panel: melting curves for adenine containing oligo in 200mM Tris-HCl 100mM $MgCl_2$ pH 8. Bottom left panel: melting curves for adenine containing oligos in 5mM Tris HCl 1M NaCl pH 7. Right panel: Van't Hoff plot for free energy determination. Continuous lines represent prediction interval for the linear fit with 0.95 confidence level for five independent data points of each series. Color code is the same as left panels.

### 2. CONCENTRATION OF 2-AMINOPURINE CONTAINING OLIGORIBONUCLEOTIDES

Since there are no published values for the extinction coefficients of 2Ap-containing RNA oligoribonucleotides, we tested whether the values calculated using the parameters from Xu and Nordlund (2) and IDT oligo analyzer for 2Ap-DNA were also accurate for our molecules. To do so, we performed a series of binding experiments titrating a probe RNA sequence containing 2Ap with a target complementary sequence of known concentration (determined through UV-Vis absorbance).

In the regime of [Target] >> $K_D$, every molecule of the Target will immediately bind to the probe, quenching the 2Ap signal in a linear fashion. This process goes on until the probe is completely saturated, switching to a Target-independent signal. The transition between these two regimes marks the true concentration of the Probe. The oligonucleotide extinction coefficient at this point can be simply calculated by dividing the absorbance measured at 260nm in a 1cm cuvette for the probe concentration determined by titration.

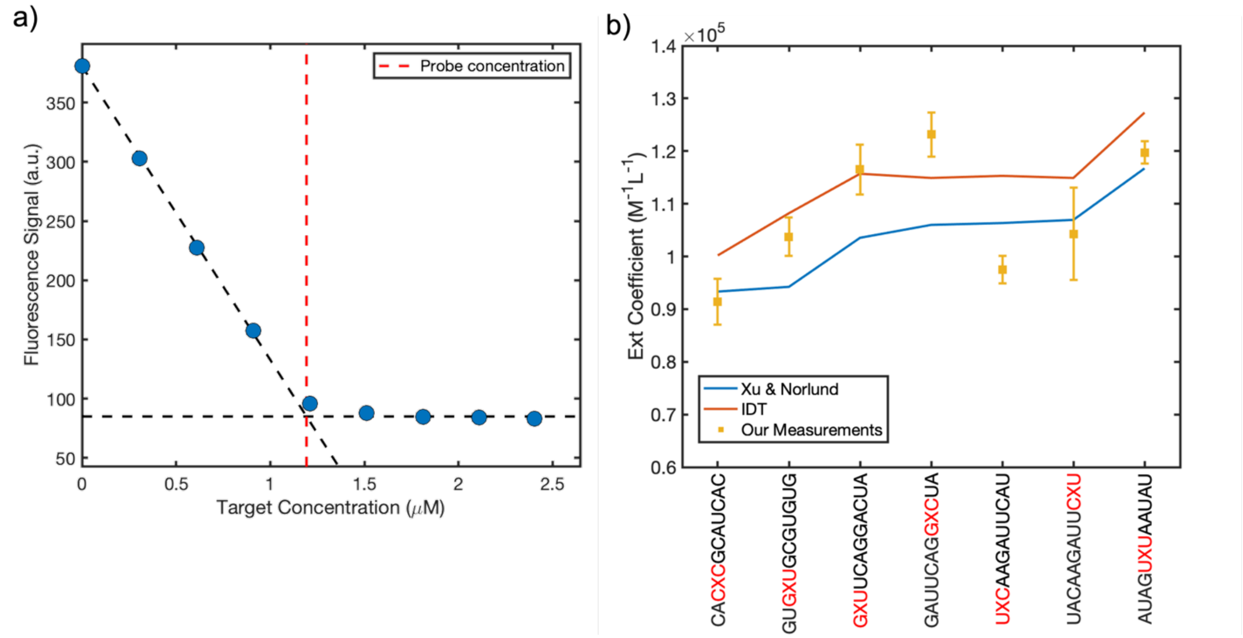

**Supplementary Figure 2.** (a) Example of extinction coefficient determination for a 2Ap-containing RNA oligonucleotide (probe) titrated with a complementary target. The transition between linear behavior and saturating regime marks the concentration of the probe. (b) Comparison between experimentally determined extinction coefficients and the ones predicted using Xu & Nordlund parameters or IDT oligo analyzer.

### 3. ANALYSIS OF STOPPED-FLOW DATA

Measurements of association kinetics were performed using a Jasco FP-8500 Spectrofluorometer equipped with the SFS-852T Stopped-Flow accessory monitoring the fluorescence emission of a 2Ap-containing oligonucleotide.

Settings used for acquisitions are the following:

- Ex bandwidth 5 nm
- Em bandwidth 10 nm
- Ex wavelength 300 nm
- Em wavelength 370 nm
- Flow time 40 ms
- Mixing ratio 1:1 using 5 ml syringes (200 μl total volume per measurements)

For each annealing reaction between the fluorescent oligonucleotide (A) and the non-fluorescent complementary oligonucleotide (B) we measure a fluorescent signal related to the variation of [A] over time. Each fluorescent trace measured this way ($F$) is converted to concentration [A] using the fluorescence value reached at complete binding ($F_\infty$) and a reference constant fluorescent trace ($F_{\mathrm{control}}$) produced mixing A with buffer:

$$[A](t) = (F(t) - F_\infty)/(F_{\mathrm{control}} - F_\infty) \cdot [A]_0$$

For long oligonucleotides studied in this work the reaction goes to completion in roughly one second with complete binding. In this regime our data are only sensitive to an upper limit of $k_{off}$ since the dissociation rate is fundamentally negligible. We can thus fit our data [A](t) with a simple second order reaction and extract $k_{on}$. As a rule of thumb, the data are sensitive to $k_{off}$ when it is roughly comparable to $k_{on}[B]_0 10^{-2}$, which corresponds to $\approx 10^{-1} s^{-1}$ and a binding affinity $K_D$ equal to $10^{-8}$ M. For short oligonucleotides having $K_D > 10^{-8}$ M ($\Delta$G > - 11 kcal/mol), dissociation is not negligible, and binding is not complete in the micromolar range. In this case a series of measurements at increasing concentrations of oligonucleotide B have been fit altogether with shared $k_{on}$ and $k_{off}$:

1) $d[A]/dt = - [A]\cdot[B]\cdot k_{on} + [AB]\cdot k_{off}$
2) $d[B]/dt = - [A]\cdot[B]\cdot k_{on} + [AB]\cdot k_{off}$
3) $d[AB]/dt = [A]\cdot[B]\cdot k_{on} - [AB]\cdot k_{off}$

For such cases, the MATLAB *fminsearch* function has been employed to minimize the residuals from the experimentally measured [A](t) and that calculated from the set of differential equations solved with *ODE15s*. The best fit parameters error has been determined from the sum square error as described in literature (3) using a threshold calibrated on our dataset as equal to 0.9.

### 3. COMPARISON WITH NN CALCULATIONS

Once we determined that the 2Ap effect on RNA duplex stability at room temperature is negligible, we asked how our measurements compare with predictions from the NN database. We gathered $\Delta$G values from multiple techniques, finding in every case a good agreement with NUPACK predictions as shown in Supplementary Figure 3.

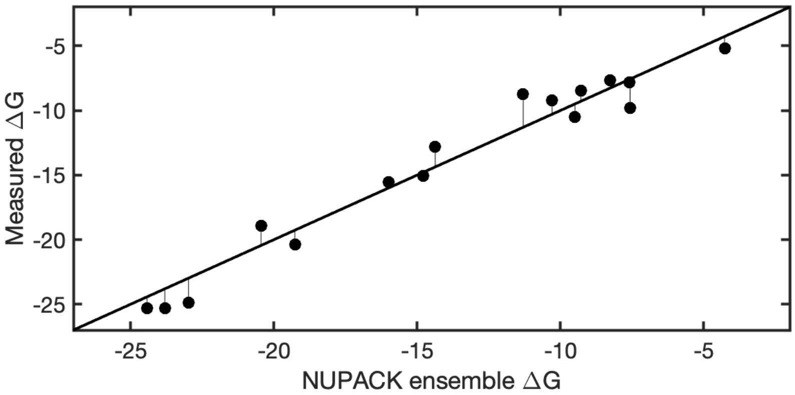

**Supplementary Figure 3.** Comparison of measured and predicted $\Delta$G at 25°C.

### 4. COMPARISON OF UV MELTING AND FLUORESCENCE MELTING

We decided to test whether some discrepancy from the NN-determined $\Delta$G at room temperature could be due to differences between fluorescence melting and UV melting results. To evaluate this, we followed the melting of a 2Ap oligonucleotide through fluorescence while changing temperature and compared it to the regular adenine-containing oligonucleotide measured through UV melting.
Results are shown below for sequences 12a (or 12a*) and 8b, and suggest that no methodological difference is present.

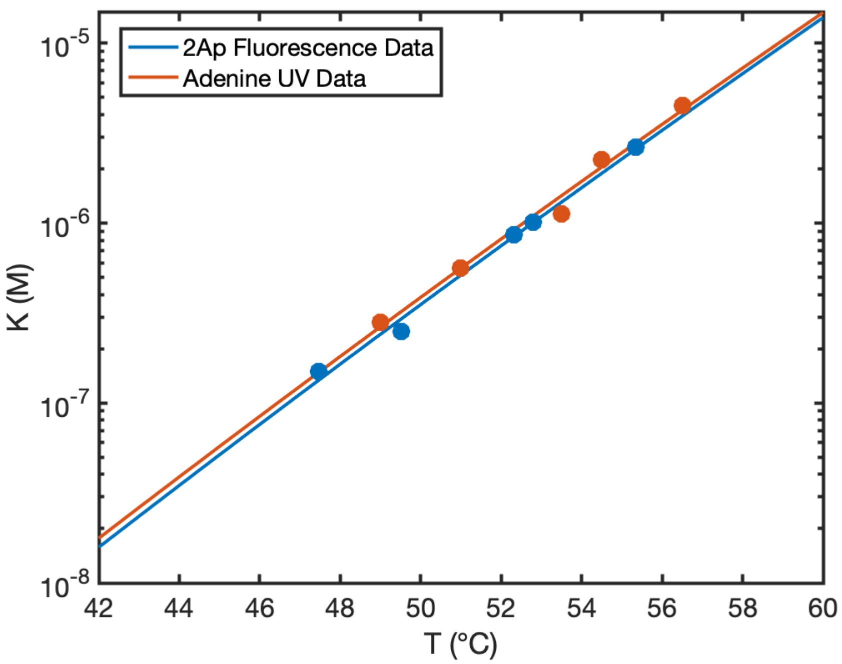

**Supplementary Figure 4.** Comparison of melting temperatures measured tracking either fluorescence emission (blue) in a 2Ap-containing oligonucleotide or UV absorbance (orange) in a regular adenine-containing oligonucleotide with the same sequence.

### 5. COMPARISON OF BUFFER CONDITIONS

To check whether our main buffer condition (100mM Tris Buffer pH 8.0 200mM $MgCl_2$) could produce results comparable with a more standard condition (5mM Tris Buffer pH 7.0 1M NaCl) we performed a subset of measurements in the latter buffer. In Supplementary Section 1 we show that the thermodynamics is not affected. Here we show a global fit for the hybridization of a 8nt long oligo that clearly shows how $k_{on}$, $k_{off}$ and $K_D$ are identical. Oligonucleotides used are A: UACAAGAUUC2ApU and B: AUGAAUCU.

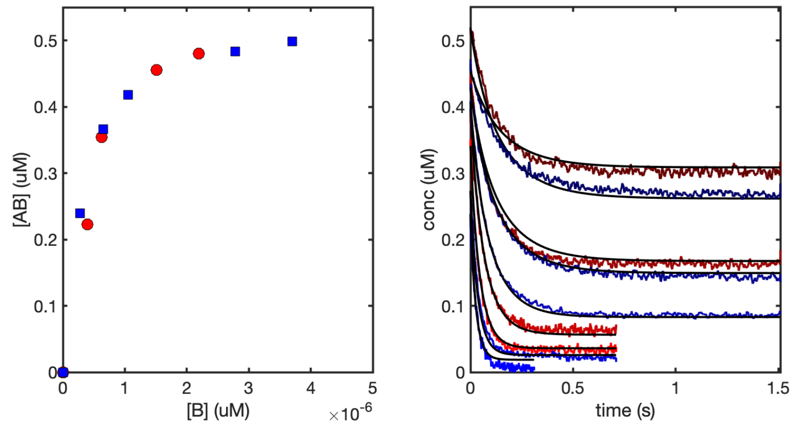

**Supplementary Figure 5** Left panel: binding curves for A and B oligonucleotides in either 100mM Tris-HCl Buffer pH 8.0 200mM $MgCl_2$ (red) or 5mM Tris-HCl Buffer pH 7.0 1M NaCl (blue) are undistinguishable. Right panel: global fit for two independent datasets of hybridization reactions in two buffer conditions show that the two sets of kinetic parameters are the same. Color code is the same as left panel.

Parameters obtained through this characterization are $k_{on}$ 1.09e7 $M^{-1}s^{-1}$, $k_{off}$ 1.35$s^{-1}$ and $K_D$ 1.24e-7 $M^{-1}$.

### 6. FOUR-WAY STRAND EXCHANGE

To rule out the contribution of four-way strand exchange in our studies, we prepared 4 equimolar mixtures of pre-annealed duplexes (8a-12b* and 8b-12a) with 4nt-long complementary overhangs, at different total concentrations. In all these mixtures the expected concentration of free single stranded oligonucleotides during the reaction varies by less than a factor of two, while the total concentration of duplexes varies by a factor of 60. If an equilibration pathway going through the direct interaction of the two duplexes does exist, we would expect to see a concentration-dependence of our time traces. The concentration-independence of the measured time traces points at a negligible four-way strand exchange, with an upper limit for the bimolecular rate approximately equal to $10^2$ $M^{-1}s^{-1}$, a value that would otherwise lead to a measurable acceleration of our reactions.

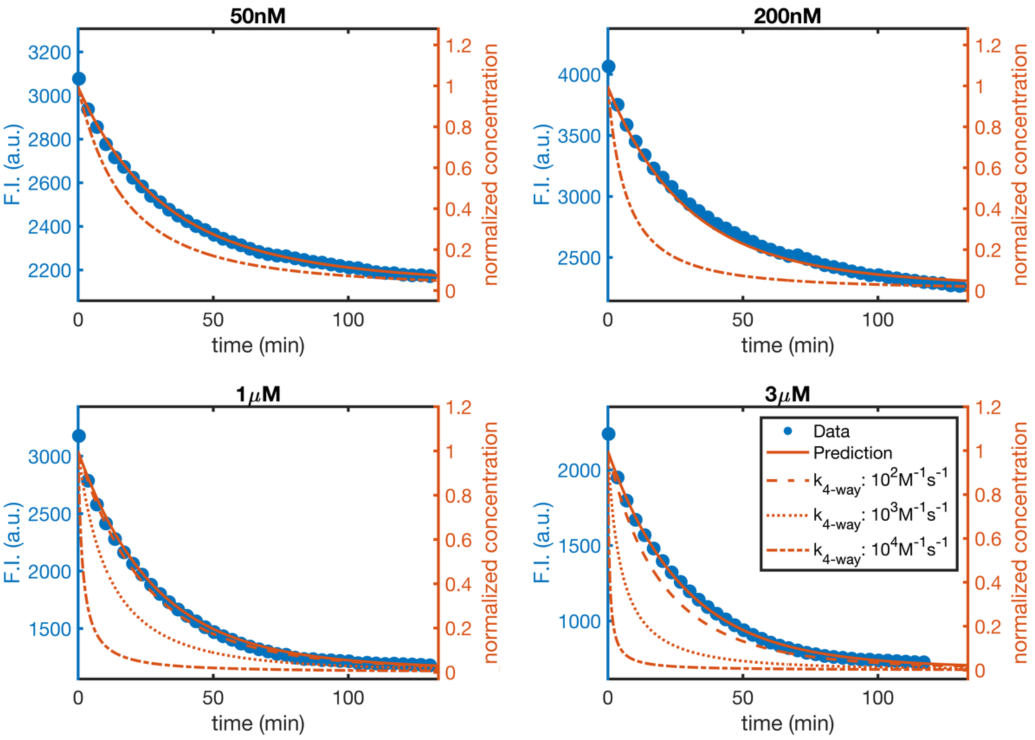

**Supplementary Figure 6** Time traces for the equilibration of mixtures of pre-annealed duplexes exposing complementary overhangs. The signal of 2Ap in the 12b* overhang gets quenched over time. The concentration written on each panel is for the two duplexes mixed at equal concentration. All time traces follow the same time-course, showing no significant concentration-dependence. Orange lines show predictions without (continuous line) and with (dotted and dashed) strand exchange having different bimolecular rates.

### 7. PREDICTION OF ANNEALING IN MIXTURES

To determine the evolution over time of a mixture of RNA oligonucleotides, we define a system of differential equations to describe the behavior of each component. For a symmetric system as the one studied in this work, where the length and sequence of the bound stretches in $A_LB_S$, $A_SB_L$ and $A_SB_S$ is the same, we can describe the reactions occurring as follows (association phenomena are highlighted in blue and strand displacement processes are highlighted in red for better clarity):

1) $[A_L]/dt = -[A_L]\cdot[B_L]\cdot k^L_{on} - [A_L]\cdot[B_S]\cdot k^S_{on} + [A_LB_L]\cdot k^L_{off} + [A_LB_S]\cdot k^S_{off} - [A_SB_L]\cdot[A_L]\cdot k_{displ}$
2) $[A_S]/dt = -[A_S]\cdot[B_L]\cdot k^L_{on} - [A_S]\cdot[B_S]\cdot k^S_{on} + [A_SB_L]\cdot k^S_{off} + [A_SB_S]\cdot k^S_{off} + [A_SB_L]\cdot[A_L]\cdot k_{displ}$
3) $[B_L]/dt = -[A_L]\cdot[B_L]\cdot k^L_{on} - [A_S]\cdot[B_L]\cdot k^S_{on} + [A_LB_L]\cdot k^L_{off} + [A_SB_L]\cdot k^S_{off} - [A_LB_S]\cdot[B_L]\cdot k_{displ}$
4) $[B_S]/dt = -[A_L]\cdot[B_S]\cdot k^S_{on} - [A_S]\cdot[B_S]\cdot k^S_{on} + [A_LB_S]\cdot k^S_{off} + [A_SB_S]\cdot k^S_{off} + [A_LB_S]\cdot[B_L]\cdot k_{displ}$
5) $[A_LB_L]/dt = [A_L]\cdot[B_L]\cdot k^L_{on} - [A_LB_L]\cdot k^L_{off} + [A_LB_S]\cdot[B_L]\cdot k_{displ} + [A_SB_L]\cdot[A_L]\cdot k_{displ}$

6) $[A_LB_S]/dt = [A_L]\cdot[B_S]\cdot k^S_{on} - [A_LB_S]\cdot k^S_{off} - [A_LB_S]\cdot[B_L]\cdot k_{displ}$
7) $[A_SB_L]/dt = [A_S]\cdot[B_L]\cdot k^L_{on} - [A_SB_L]\cdot k^S_{off} - [A_SB_L]\cdot[A_L]\cdot k_{displ}$
8) $[A_SB_S]/dt = [A_S]\cdot[B_S]\cdot k^S_{on} - [A_SB_S]\cdot k^S_{off}$

All parameters have been calculated as described in the main text:
- $k_{displ}$ is computed as $k_s \cdot K_A$, with $k_s$ = 12.4 $s^{-1}$. Two displacement rates should be considered here, one for $A_L$ displacing $A_SB_L$ and one for $B_L$ displacing $A_LB_S$. For a symmetric system the two numbers are going to be extremely close and can be approximated as a single rate.
- Association kinetics $k^L_{on}$ and $k^S_{on}$ have been calculated using Eq.10 and Eq. 11 from main text.
- Dissociation kinetics $k^L_{off}$ and $k^S_{off}$ have been calculated as $k_{on} \cdot e^{\Delta G/(R.T),}$ where ΔG is the binding energy of the two associated oligonucleotides.

To determine a prediction interval, we varied the parameters within their typical experimental error found in the context of this work: 3% for ΔG, 10% for $k_{on}$ and 5% for $\Delta G_{toehold}$.

Sequences used in the experiments are the following (see Supplementary Data 8):

| | $A_S$ | $B_S$ | $A_L$ | $B_L$ |
|---|---|---|---|---|
| Mix a) | 8a | 8b | 12a | 12b* |
| Mix b) | 8a | 8b | 11a | 12b* |
| Mix c) | 8a | 8b | 10a | 12b** |
| Mix d) | 8a | 8b | 10a | 12b** |

### 8. LIST OF SEQUENCES USED

| Sequence name | Sequence | CG content | Template Code |
|---|---|---|---|
| 12a | GUG AUG CGU GUG | 0.58 | V |
| 12a* | GUG XUG CGU GUG | 0.58 | V |
| 12b | CAC ACG CAU CAC | 0.58 | V |
| 12b** | CXC ACG CAU CAC | 0.58 | V |
| 12b* | CAC XCG CAU CAC | 0.58 | V |
| 11a | GUG AUG CGU GU | 0.55 | V |
| 10a* | GUG XUG CGU G | 0.60 | V |
| 10a | GUG AUG CGU G | 0.60 | V |
| 10b | C ACG CAU CAC | 0.60 | V |
| 9a | GUG AUG CGU | 0.56 | V |
| 8a | GUG AUG CG | 0.63 | V |
| 8b | CG CAU CAC | 0.63 | V |
| 7a | GUG AUG C | 0.57 | V |
| 6b | CAU CAC | 0.50 | V |
| T* | CXC GCA UCA CCA | 0.58 | T |
| T | CAC GCA UCA CCA | 0.58 | T |
| T12 | UGG UGA UGC GUG | 0.58 | T |
| T11 | GG UGA UGC GUG | 0.64 | T |
| T10 | G UGA UGC GUG | 0.60 | T |
| T9 | UGA UGC GUG | 0.56 | T |
| T8 | GA UGC GUG | 0.63 | T |
| T7 | A UGC GUG | 0.57 | T |
| T6 | UGC GUG | 0.67 | T |
| T5 | GC GUG | 0.80 | T |
| T4 | C GUG | 0.75 | T |
| MC* | GAU UCA GGX CUA | 0.42 | M |
| MA* | GXU UCA GGA CUA | 0.42 | M |
| M12 | UAG UCC UGA AUC | 0.42 | M |
| M11 | UAG UCC UGA AU | 0.36 | M |
| M10 | UAG UCC UGA A | 0.40 | M |
| M9 | UAG UCC UGA | 0.44 | M |
| M8 | UAG UCC UG | 0.50 | M |
| M7 | UAG UCC U | 0.43 | M |
| K* | UAC AAG AUU CXU | 0.25 | K |
| K12 | AUG AAU CUU GUA | 0.25 | K |
| K10 | AUG AAU CUU G | 0.30 | K |
| K9 | AUG AAU CUU | 0.22 | K |
| K8 | AUG AAU CU | 0.25 | K |
| S* | AUAGUXUAAUAU | 0.08 | S |
| S12 | AUAUUAUACUAU | 0.08 | S |
| S11 | AUAUUAUACUA | 0.09 | S |
| A* | AAUAAAUAAAXU | 0.00 | A |
| A12 | AUUUUAUUUAUU | 0.00 | A |

Note: **X** refers to adenine to 2-aminopurine substitution.

### 9. LIST OF BINDING EXPERIMENTS PERFORMED

| Sequence 1 | Sequence 2 | $k_{on}$ ($M^{-1}s^{-1}$) | std |
|---|---|---|---|
| T* | T12 | 3.25E+07 | 4.46E+06 |
| T* | T11 | 3.50E+07 | 4.62E+06 |
| T* | T10 | 2.84E+07 | 4.26E+06 |
| T* | T9 | 2.14E+07 | 3.58E+06 |
| T* | T8 | 2.64E+07 | 5.06E+06 |
| T* | T7 | 1.61E+07 | 2.35E+06 |
| T* | T6 | 1.79E+07 | 1.56E+06 |
| T* | T5 | 1.59E+07 | 2.33E+05 |
| MC* | M12 | 1.86E+07 | 2.25E+06 |
| MC* | M11 | 1.58E+07 | 1.70E+06 |
| MC* | M10 | 1.64E+07 | 1.37E+06 |
| MC* | M9 | 1.54E+07 | 1.31E+06 |
| MC* | M8 | 1.35E+07 | 1.25E+06 |
| MC* | M7 | 7.55E+06 | 9.82E+05 |
| K* | K12 | 1.51E+07 | 2.46E+06 |
| K* | K10 | 1.19E+07 | 5.61E+05 |
| K* | K9 | 1.09E+07 | 5.87E+05 |
| K* | K8 | 1.17E+07 | 5.84E+05 |
| 12a* | 12b | 3.83E+07 | 2.02E+06 |
| 12a* | 8b | 2.14E+07 | 3.30E+06 |
| 12a* | 6b | 6.53E+06 | 5.25E+05 |
| S* | S12 | 7.93E+06 | 3.65E+05 |
| S* | S11 | 7.37E+06 | 4.95E+05 |
| A* | A12 | 1.39E+06 | 6.19E+04 |

### 10. LIST OF STRAND DISPLACEMENT EXPERIMENTS PERFORMED

| Oligo B | Oligo A | Oligo C | $k_{displ}$ ($M^{-1}s^{-1}$) | std | N | ΔG (kcal/mol) | toehold |
|---|---|---|---|---|---|---|---|
| 12b* | 8a | 11a | 3.68E+04 | 8.50E+02 | 3 | 4.56 | 3nt (5') |
| 12b* | 9a | 10a | 6.42E+02 | 3.96E+01 | 2 | 1.05 | 1nt (5') |
| T | 10a* | 11T | 9.04E+02 | 4.79E+01 | 3 | 3.05 | 1nt (3') |
| 12b** | 9a | 12a | 6.45E+04 | 5.87E+03 | 3 | 5.36 | 3nt (5') |
| 12b* | 8a | 10a | 2.75E+03 | 3.54E+02 | 3 | 2.8 | 2nt (5') |
| 12b | 10a* | 10a | 5.53E+00 | 1.88E+00 | 4 | *-1.68*# | 0 |
| 12a | 12b* | 12b | 2.05E+00 | 2.18E-01 | 3 | *-1.68*# | 0 |
| 12b** | 8a | 12a | 1.73E+06 | 9.19E+04 | 3 | 7.12 | 4nt (5') |
| 12b** | 10a | 12a | 1.53E+02 | 1.41E+01 | 3 | 2.03 | 2nt (5') |
| 12a | 8b | 12b** | 8.25E+05 | 3.61E+04 | 3 | 7.02 | 4nt (3') |
| 12a | 10b | 12b** | 2.44E+03 | 4.81E+01 | 3 | 1.93 | 2nt (3') |
| MA | M10 | M12 | 1.74E+02 | 1.44E+01 | 3 | 1.56 | 2nt (5') |
| MA | M8 | M12 | 1.25E+06 | 2.17E+05 | 3 | 5.56 | 4nt (5') |
| MA | M9 | M12 | 1.81E+03 | 1.40E+02 | 3 | 2.86 | 3nt (5') |
| 12b** | 7a | 12a | 2.71E+06 | 5.26E+05 | 3 | 10.71 | 5nt (5') |
| 12b** | 7a | 11a | 2.04E+06 | 5.40E+05 | 3 | 8.15 | 4nt (5') |

#Apparent toehold binding energies calculated as described in main text.
